# Copper sensing by OXA1L triggers a mitochondria-to-nucleus stress response to restrain cuproptosis

**DOI:** 10.64898/2026.11.17.767466

**Authors:** Xueliang Yang, Chong Wei, Ye Lu, Qinglin Li, Xin Liu, Gang Shao, Shuyuan Zhang, Hengzhi Li, Yuzhe Wang, Keke Zheng, Xiaoyan Dai, Jian Kang, Chulin Sha, Qiang Zhang, Caiyun Fu

## Abstract

Copper overload triggers cuproptosis, a regulated cell death driven by aggregation of lipoylated DLAT and collapse of mitochondrial proteostasis. How cells sense this stress and mount an adaptive response is unknown. Here, combining time-resolved proximity-labeling proteomics across five subcellular compartments with genetic and biochemical dissection, we define a mitochondria-to-nucleus retrograde surveillance pathway. Copper binding to the inner-membrane insertase OXA1L requires His170, His172 and His209, disrupting OXA1L function and MRPL20 incorporation into the assembling 39S mitoribosome. Processed MRPL20 subsequently translocates to the nucleus, where it binds to ATF5 and drives activation of the mitochondrial unfolded protein response (UPR^mt^). Parallelly, the integrated stress response (ISR) converges on ATF5 to further amplify UPR^mt^ output via transcriptional and translational upregulation of ATF5 abundance. Functionally, UPR^mt^ activation limits DLAT oligomerization and restrains cuproptosis. Pharmacological inhibition of the UPR^mt^ enhances copper-induced cytotoxicity and suppresses xenograft growth, uncovering a therapeutic vulnerability for copper-based cancer therapy.

## Main Text

Copper is an essential cofactor for numerous cellular enzymes, yet excess intracellular copper can disrupt protein homeostasis and contribute to diverse human diseases (1, 2). The discovery of cuproptosis established a direct link between mitochondrial copper accumulation and a distinct form of regulated cell death characterized by aggregation of the lipoylated tricarboxylic acid cycle proteins, including DLAT, loss of Fe-S cluster proteins, and collapse of mitochondrial proteostasis (3). These features raise a fundamental question: how do cells sense copper-triggered mitochondrial proteotoxic stress and what endogenous mechanisms restrain its progression to cell death?

The mitochondrial unfolded protein response (UPR^mt^) is a conserved transcriptional surveillance program activated by mitochondrial proteotoxic stress, functioning to restore proteostasis by transcriptionally upregulating mitochondrial chaperones, proteases and other adaptive factors (4). Because toxic protein aggregation is central to cuproptosis, we hypothesized that the UPR^mt^ may provide an endogenous defense against copper-induced proteotoxicity. However, how mitochondrial copper excess is sensed and coupled to UPR^mt^ activation remains unclear, and conventional static proteomic approaches are not suited to resolve the dynamic inter-organelle protein redistribution that relays mitochondrial stress signals to the nucleus. Here we identify the inner-membrane insertase OXA1L as a mitochondrial copper sensor and define an OXA1L-MRPL20-ATF5 retrograde signaling pathway that activates UPR^mt^ and limits cuproptosis.

### Spatio-temporal proteomic profiling reveals compartment-specific remodeling during early cuproptosis

To characterize subcellular changes accompanying early copper stress, we exposed HT-1080 fibrosarcoma cells to the copper ionophore elesclomol plus CuSO (ES-Cu) and examined cellular ultrastructure by transmission electron microscopy (TEM) across early time points (Fig. S1A–S1C). At 12 hours, mitochondria were enlarged but retained a largely intact, electron-dense matrix and partially preserved cristae architecture. By 24 hours, mitochondria exhibited severe swelling, cristae fragmentation and extensive vacuolization, accompanied by abnormalities of the nuclear envelope and fragmentation of the endoplasmic reticulum (ER) into small vesicle-like structures. No overt plasma membrane rupture or apoptotic body formation was detected. These morphological changes indicated progressive remodeling of multiple organelles during copper stress and motivated a dynamic analysis of subcellular proteome organization.

To capture spatial and temporal proteome changes during this early response, we developed a miniTurbo-based proximity-labeling mass spectrometry strategy (5) termed Spatio-Temporal Mass Spectrometry (STMS). MiniTurbo reporters were targeted to five core subcellular compartments, including the ER, Golgi apparatus, nucleus, outer mitochondrial membrane (OMM) and plasma membrane (PM) (6), to capture compartment-proximal proteomes over time (Fig.1A). Immunofluorescence and immunoblotting confirmed the expected localization of each miniTurbo reporter under basal conditions and after ES-Cu treatment (Fig. 1B and fig.S2A), supporting the spatial fidelity of the labeling system.

**Fig. 1.**
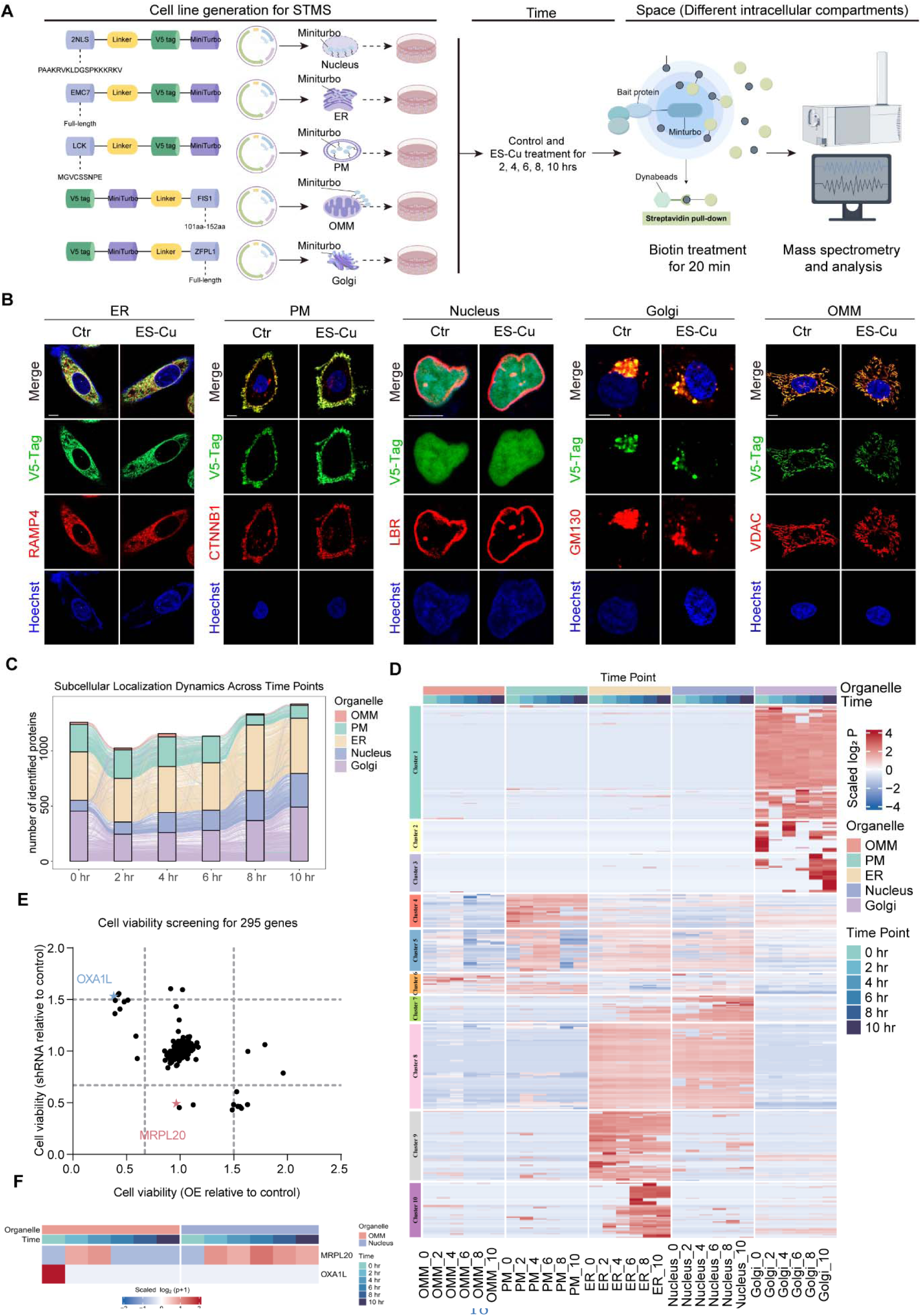
Spatio-temporal proteomics of early cuproptosis nominates OXA1L and MRPL20 as candidate regulators. **(A)** Schematic of spatio-temporal mass spectrometry (STMS) workflow. HT-1080 cells expressing compartment-targeted miniTurbo were exposed to elesclomol–copper (ES-Cu; 15 nM elesclomol and 1.5 μM CuSO ) for 0, 2, 4, 6, 8 or 10 h, biotin-labeled for 20 min, and processed for streptavidin enrichment and mass spectrometry. The reporters targeted the endoplasmic reticulum (ER), plasma membrane (PM), nucleus, outer mitochondrial membrane (OMM) and Golgi apparatus. **(B)** Representative immunofluorescence images validating the subcellular distribution of the five miniTurbo reporters. V5-tagged reporters (green) are shown with the indicated organelle markers (red) under control and ES-Cu conditions. Nuclei were counterstained with Hoechst (blue). Scale bars, 10 μm. **(C)** Alluvial plot of changes in the dominant subcellular assignment of identified proteins across the six sampling times. Stacked bars show the numbers assigned to each organelle, and connecting flows represent proteins changing their predominant localization. **(D)** Heatmap of normalized protein abundance across the five labeled compartments and the six sampling times. Rows are grouped into 10 spatio-temporal clusters; color indicates scaled log protein abundance. **(E)** Functional screen of 295 candidate genes under ES-Cu treatment. Each point represents a candidate; the x and y axes show cell viability relative to the corresponding controls after overexpression (OE) and shRNA knockdown, respectively. OXA1L and MRPL20 are highlighted. **(F)** STMS-derived compartment and time profiles of MRPL20 and OXA1L. The heatmap summarizes the relative abundance of each protein in the indicated compartments and sampling times.

We collected proximity-labeled proteomic samples at baseline and five sequential time points after ES-Cu exposure to track early proteome remodeling, and identified more than 4,000 proteins, of which 2,867 passed stringent quality filtering (Table S1). Biological replicates showed high reproducibility and distinct compartment-associated proteomic profiles (fig. S2B and S2C). The numbers of quantified proteins in the streptavidin-enriched fraction rose sharply at the PM and OMM within the first 2 hours and then fell to below half of initial levels by 6 or 8 hours, whereas the ER and nucleus increased steadily, and the Golgi showed temporal fluctuations (fig. S2D). In contrast, summed protein intensity within the biotinylated pool showed a different pattern, including a progressive decline in the ER with smaller overall changes in the other compartments (fig. S2E). These readouts describe the enriched proximity-labeled fraction rather than total organelle protein content. Analysis of proteins whose proximity-labeling abundance changed significantly relative to baseline further revealed distinct compartment-specific temporal responses. The nucleus and ER showed a progressive increase in the number of upregulated proteins, whereas the OMM and PM exhibited biphasic patterns, with an early surge in upregulated proteins followed by a later increase in downregulated proteins (fig S3A-S3E, table S2).

We next inferred potential protein relocalization by assigning each protein to its dominant compartment using a confidence scoring system benchmarked against public subcellular localization databases (7) (fig. S3F–S3H and table S3). Proteins with high confidence scores and compartment-specific localization were selected for subcellular trafficking analysis. Approximately 80% of detected proteins retained stable compartmental residency across all time points, while a defined subset changed dominant assignments between compartments, consistent with stress-induced protein redistribution (Fig. 1C).

To further characterize proteome dynamics, we categorized the subcellular proteome into 10 clusters based on their spatiotemporal expression patterns (Fig. 1D). Clusters 1, 2, 3, 9 and 10 exhibited compartment-restricted profiles, whereas the remaining clusters contained proteins distributed across multiple compartments. Most clusters displayed noticeable expression changes in response to ES-Cu treatment over time and were enriched with proteins associated with distinct molecular functions (Fig. 1D and fig. S2F), linking copper stress to activation of the ER ubiquitin-proteasome system, destabilization of Fe-S cluster proteins and disruption of translation. Detailed cluster annotations are provided in the supplementary text. Together, these analyses reveal remodeling of subcellular protein organization during early copper stress and identify a discrete set of proteins undergoing pronounced compartment-associated changes for subsequent functional investigation.

### Copper induces selective redistribution of MRPL20 from mitochondria to the nucleus

We reasoned that proteins whose abundance or subcellular distribution changed markedly under copper stress would contribute functionally to cuproptosis. We therefore performed a focused genetic screen of 295 candidates (276 top-ranked STMS hits plus 19 literature-curated proteins). shRNA-mediated knockdown and overexpression identified 28 genes whose perturbation altered cell viability by more than 30% following ES-Cu treatment (Fig. 1E and table S4). Among these hits, OXA1L and MRPL20 were prioritized because they both linked to mitochondrial protein biogenesis, and showed reciprocal effects on ES-Cu sensitivity, suggesting a functional connection (Fig. 1E). OXA1L mediates insertion of mitochondrial- and nuclear-encoded proteins into the inner mitochondrial membrane (8) and its matrix-exposed C-terminal domain interacts with proteins of the 39S mitochondrial ribosomal subunit, including MRPL20 (9, 10). STMS showed distinct but coordinated changes with a rapid decline of mitochondrial OXA1L, coincident with transient enrichment of MRPL20 at the OMM followed by its nuclear accumulation (Fig. 1F and fig. S2G).

Biochemical fractionation confirmed loss of both proteins from the mitochondrial fraction 4 hours after ES-Cu and a concomitant rise in nuclear MRPL20 (Fig. 2A). In HT-1080 cells stably expressing GFP-tagged MRPL20, live-cell confocal imaging showed exclusive mitochondrial localization at baseline. Within 4 to 10 hours of ES-Cu, the mitochondrial signal intensity rapidly declined while nuclear signal increased, with only a modest fall in total GFP intensity (Fig. 2B and 2C). Higher ionophore concentration accelerated this mitochondria-to-nucleus trafficking (fig. S4A and S4B). Immunofluorescent staining of endogenous MRPL20 recapitulated these dynamics, which was prevented by the copper chelator tetrathiomolybdate (TTM) (fig. S4C and S4D).

**Fig. 2.**
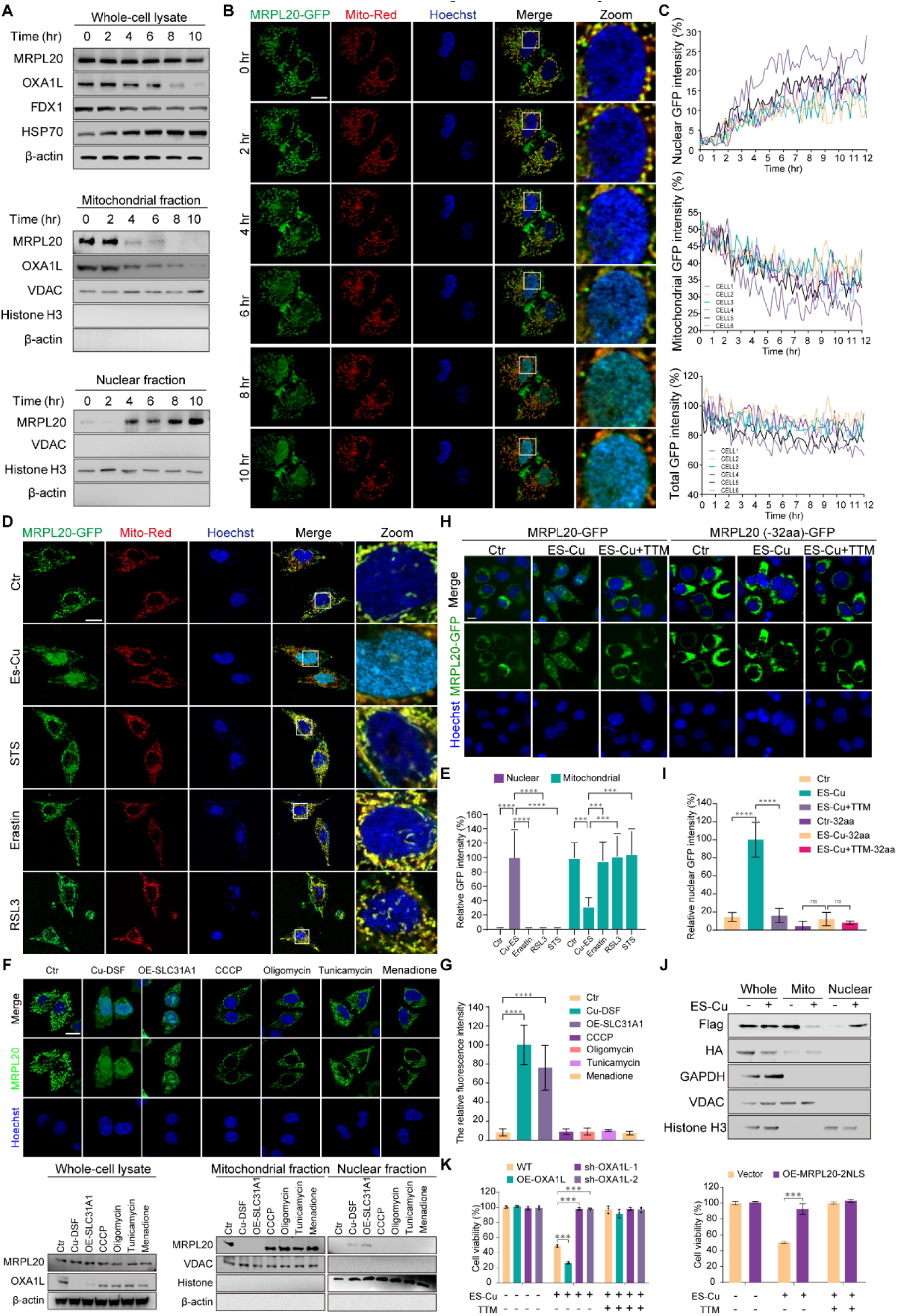
Copper induces selective redistribution of MRPL20 from mitochondria to the nucleus. **(A)** Immunoblot analysis of whole-cell lysates and mitochondrial and nuclear fractions of HT-1080 cells at the indicated times after ES-Cu treatment. MRPL20 and OXA1L were detected alongside the indicated controls; VDAC and histone H3 assess mitochondrial and nuclear fractionation, respectively. **(B)** Representative time-course confocal images of MRPL20– GFP (green), Mito-Red (red) and Hoechst (blue) in HT-1080 cells after ES-Cu treatment. The rightmost column shows enlarged nuclear regions. Scale bars, 10 μm. **(C)** Single-cell trajectories of nuclear, mitochondrial and whole-cell GFP fluorescence, expressed relative to each cell’s initial signal, during 12 h of imaging following ES-Cu exposure. **(D)** Representative MRPL20– GFP fluorescence images under vehicle, ES-Cu, staurosporine (STS), erastin or RSL3 treatment; Mito-Red and Hoechst delineate mitochondria and nuclei. Cells were treated with ES-Cu (1.5 μM CuSO plus 15 nM elesclomol, 10 h), STS (200 nM, 8 h), erastin (50 μM, 24 h), or RSL3 (200 nM, 8 h). Scale bars, 10 μm. **(E)** Quantification of nuclear and mitochondrial MRPL20– GFP fluorescence for the conditions shown in **(D)**. **(F)** Representative MRPL20 immunofluorescence images and immunoblots of whole-cell, mitochondrial and nuclear fractions after the indicated copper-overload or non-copper stress treatments. Cu-DSF denotes copper plus disulfiram; OE-SLC31A1 denotes SLC31A1 overexpression with copper. CCCP, oligomycin, tunicamycin and menadione served as non-copper stress comparators. **(G)** Quantification of nuclear MRPL20 fluorescence for the treatments shown in **(F)**. **(H)** MRPL20–GFP localization in cells expressing full-length MRPL20 or the indicated 32-amino-acid deletion mutant under control, ES-Cu or ES-Cu plus the copper chelator tetrathiomolybdate (TTM). Hoechst marks nuclei. **(I)** Quantification of the nuclear GFP signal in the conditions shown in **(H)**. **(J)** Immunoblot analysis of whole-cell, mitochondrial and nuclear fractions from cells expressing MRPL20 with an N-terminal HA tag and a C-terminal Flag tag, with or without ES-Cu. The fractionation markers GAPDH, VDAC and histone H3 are shown. **(K)** Cell viability under the indicated control, ES-Cu (1.5 μM CuSO plus 15 nM elesclomol, 10 h) or ES-Cu plus TTM conditions after OXA1L overexpression or knockdown (left), or enforced nuclear localization of MRPL20 (2NLS-MRPL20; right).

MRPL20 redistribution was copper-specific rather than a generic stress response. MRPL20 nuclear accumulation was not detected after treatment with the apoptosis inducer staurosporine or ferroptosis inducers erastin and RSL3 (11) (Fig. 2D and 2E), nor after mitochondrial uncoupling with CCCP, ATP synthase inhibition with oligomycin, oxidative stress induced by menadione, or ER stress induced by tunicamycin (Fig. 2F and 2G). Conversely, two independent copper-overload models, including copper-disulfiram (Cu-DSF) and SLC31A1 overexpression-driven copper influx, each induced pronounced MRPL20 nuclear accumulation and mitochondrial OXA1L depletion (Fig. 2F and 2G).

cNLS Mapper (12) predicted three putative NLS motifs within MRPL20 (fig. S4E). Deletion of the highest-scoring 32-amino-acid motif (RHFRGRKNRCYRLAVRTVI-RAFVKCTKARYLK) eliminated ES-Cu-induced MRPL20 nuclear import (Fig. 2H and 2I), establishing that an intrinsic targeting sequence is required for copper-stress-induced nuclear targeting. To determine the origin of the nuclear pool, we blocked cytosolic translation with cycloheximide (CHX) before ES-Cu treatment. CHX pretreatment nearly eliminated copper-induced MRPL20 nuclear enrichment in both fluorescence imaging and subcellular fractionation assays (fig. S4F-S4G), indicating that the nuclear pool depends largely on newly synthesized MRPL20. However, inhibition of protein synthesis alone cannot determine whether MRPL20 reaches the nucleus directly from the cytosol or first enters mitochondria before being released. We therefore expressed a dual-end-tagged MRPL20 reporter bearing an N-terminal HA tag upstream of its mitochondrial targeting presequence and a C-terminal Flag tag (Fig. 2J). Flag-tagged MRPL20 was detected in whole-cell lysate and in mitochondria at baseline. After ES-Cu, mitochondrial Flag signal declined while a Flag-positive species became detectable in the nuclear fraction. The N-terminal HA tag was readily detected in whole-cell lysate but present only at low levels in the mitochondrial fraction and was not detected in the nuclear fraction under either condition. Because presequence-dependent mitochondrial import is typically followed by N-terminal cleavage by mitochondrial processing peptidase (MPP) (13), the absence of HA specifically in the Flag-positive nuclear species is consistent with MRPL20 entering mitochondria and undergoing presequence processing before its nuclear appearance.

This redistribution was selective. Other OXA1L-associated large mitoribosomal proteins, including MRPL48 and MRPL51, also showed reduced mitochondrial occupancy after ES-Cu exposure but did not accumulate detectably in nuclei (fig. S4H-S4I). To assess the integrity of the canonical mitochondrial import pathway, we monitored a doxycycline-inducible mitochondrially targeted GFP reporter (fig. S4J), whose delivery depends on the presequence-mediated TOM20–TIM23 machinery (13). Newly induced reporter fluorescence remained predominantly mitochondrial over 24 h, including the late stages at which mitochondria are severely swollen (fig. S4J). These results argue against either generalized mitochondrial ribosomal protein escape or global collapse of the presequence import pathway as the basis for MRPL20 nuclear accumulation and, instead, support selective redistribution of MRPL20 after mitochondrial import and processing.

Functionally, OXA1L knockdown or enforced nuclear localization of MRPL20 (MRPL20-2NLS) protected cells from ES-Cu-induced cytotoxicity, whereas OXA1L overexpression increased sensitivity (Fig. 2K). Notably, all these effects were abolished by TTM, underscoring copper specificity. These OXA1L-dependent phenotypes were reproduced across 293T, MEF, mouse chondrocytes, U2OS, HeLa, and HepG2 cells (fig. S5A-S5F), each of which also showed ES-Cu-induced MRPL20 nuclear translocation (fig. S5G-S5L), indicating a generalizable response to copper stress.

### Copper binding inactivates OXA1L

Given the rapid loss of OXA1L following copper influx, we asked whether OXA1L acts as a sensor of copper overload. Metalloprotein prediction (MIB2, an updated implementation of MIB) (14) identified OXA1L as a high-confidence copper-binding protein, with two predicted binding sites receiving the highest confidence score (fig. S6A). Consistent with these predictions, OXA1L from cell lysates was retained preferentially on copper-charged resin and weakly on magnesium-charged resin (Fig. 3A and 3B). Domain mapping localized copper binding to the N4 fragment (aa 163-216) (Fig. 3C and 3D), concordant with MIB2 predicted pockets (fig. S6B-S6C, Table S5). Alanine scanning across aa 109-216 pinpointed His170, His172 and His209 as essential residues for copper binding (Fig. 3E). A triple mutant (H170A/H172A/H209A; OXA1L-Mut) abolished copper-resin binding while preserving magnesium-resin binding (Fig. 3F). OXA1L-Mut was also resistant to ES-Cu-induced protein loss (Fig. 3G), linking these copper-binding residues to stress-induced OXA1L destabilization.

**Fig. 3.**
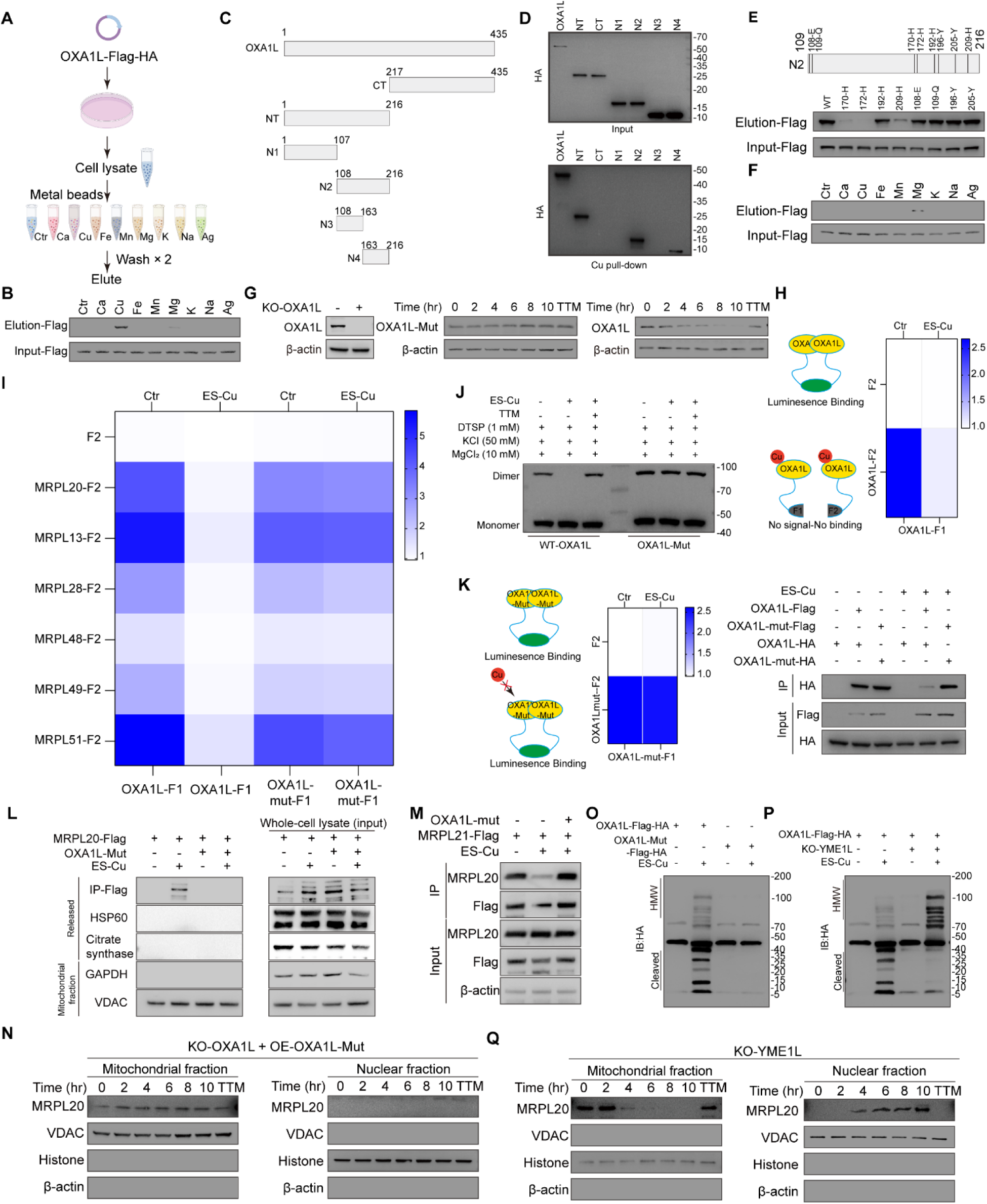
Copper binds OXA1L at H170/H172/H209 and inactivates it, releasing MRPL20. **(A)** Schematic of the metal-affinity pull-down assay using lysates containing tagged OXA1L and resins charged with the indicated metal ions. **(B)** Anti-Flag immunoblots of input lysates and metal-resin eluates from cells expressing tagged OXA1L. **(C)** Schematic of full-length OXA1L and the indicated truncations used to map its copper-binding region. **(D)** Copper-affinity pull-down of the OXA1L truncations in **(C)**, detected by anti-HA immunoblot in input and eluate fractions. **(E)** Copper-affinity pull-down of the indicated histidine-to-alanine OXA1L mutants in the 109–216 region. Anti-Flag immunoblots show eluates and inputs. **(F)** Metal-affinity pull-down of the copper-binding-deficient triple mutant H170A/H172A/H209A (OXA1L-Mut), using the metal-charged resins indicated in **(A)**. **(G)** OXA1L-knockout validation (left) and time-course immunoblots of wild-type OXA1L and OXA1L-Mut following ES-Cu treatment, with β-actin as a loading control; a TTM condition is included as indicated. **(H)** Schematic and NanoLuc complementation readout of wild-type OXA1L homodimerization under control and ES-Cu conditions. Luminescence is reported as a normalized luminescence ratio (NLR). **(I)** NanoLuc complementation heatmap of wild-type OXA1L or OXA1L-Mut interactions with the indicated 39S mitoribosomal proteins (MRPL20, MRPL13, MRPL28, MRPL48, MRPL49 and MRPL51) under control and ES-Cu conditions. **(J)** Immunoblot of monomeric and dimeric OXA1L species after DTSP cross-linking under the indicated copper, salt and magnesium conditions for wild-type OXA1L and OXA1L-Mut. **(K)** NanoLuc complementation analysis of OXA1L-Mut self-association (schematic and heatmap, left) and co-immunoprecipitation of differently tagged wild-type or mutant OXA1L under the indicated ES-Cu conditions (right). **(L)** Cell-free release assay using mitochondria isolated from cells expressing MRPL20–Flag with or without OXA1L-Mut. Isolated mitochondria were incubated with or without ES-Cu in mitochondrial storage buffer. The released fraction was analyzed for MRPL20–Flag after anti-Flag immunoprecipitation and for HSP60 and citrate synthase; GAPDH and VDAC were assessed in the mitochondrial fraction. Whole-cell lysates are shown as inputs. **(M)** Co-immunoprecipitation of MRPL21–Flag and MRPL20 under the indicated ES-Cu and OXA1L-Mut conditions, with input controls. **(N)** Immunoblot time course of mitochondrial and nuclear MRPL20 in OXA1L-knockout cells reconstituted with OXA1L-Mut following ES-Cu treatment. TTM and fractionation controls are shown. **(O)** Anti-HA immunoblot of high-molecular-weight OXA1L species and lower-molecular-weight fragments in cells expressing tagged wild-type OXA1L or OXA1L-Mut after ES-Cu treatment. **(P)** Anti-HA immunoblot of OXA1L high-molecular-weight species and fragments in control and YME1L-knockout cells after ES-Cu. **(Q)** Immunoblot time course of mitochondrial and nuclear MRPL20 in YME1L-knockout cells following ES-Cu treatment, with TTM and fractionation controls.

OXA1L functions as a homodimeric inner-membrane insertase that associates with the large mitoribosomal subunit (8, 9, 10). In NanoLuc binary complementation assays (15), ES-Cu reduced OXA1L homodimerization (Fig. 3H) and its association with large-subunit ribosomal proteins MRPL13, MRPL20, MRPL28, MRPL48, MRPL49 and MRPL51 (Fig. 3I). Chemical cross-linking independently confirmed the loss of OXA1L dimers after ES-Cu treatment (Fig. 3J). This copper-mediated disruption of OXA1L homodimerization can be prevented by mutation of copper binding sites in OXA1L (Fig. 3H–3K), establishing that direct copper binding, but not a downstream consequence of copper toxicity, inactivates OXA1L.

We next asked whether OXA1L inactivation causally drives MRPL20 redistribution during copper stress. In a cell-free assay using isolated mitochondria, ES-Cu triggered release of MRPL20-Flag, whereas expression of the copper-binding deficient OXA1L-Mut suppressed this release (Fig. 3L). ES-Cu also weakened the association between MRPL20 and MRPL21, another component of the 39S mitoribosomal subunit (16), and OXA1L-Mut preserved this interaction (Fig. 3M). Because 39S biogenesis requires coordinated assembly of mitoribosomal proteins onto mitochondrial rRNA, these observations are consistent with impaired incorporation of MRPL20 into the assembling large subunit after OXA1L inactivation. In intact cells, OXA1L knockdown was sufficient to increase nuclear MRPL20, while OXA1L overexpression suppressed ES-Cu-induced MRPL20 translocation (fig. S7A-S7D). Reconstitution of OXA1L-deficient cells with OXA1L-Mut also prevented MRPL20 appearance in the nuclear fraction (Fig. 3N). Imaging confirmed that MRPL20 remained predominantly mitochondrial under these conditions (fig. S7E). Together, these results place OXA1L functional inactivation upstream of MRPL20 mitochondrial release and nuclear accumulation.

Finally, we asked how copper stress affects OXA1L protein stability. The degradation of OXA1L was independent of the ubiquitin-proteasome and lysosomal pathways (fig. S7F) but required the IMM metalloprotease YME1L, whose knockout stabilized OXA1L (fig. S7G). Copper treatment produced high-molecular-weight (HMW) species of tagged OXA1L and lower-molecular-weight cleavage fragments, both of which were markedly reduced in OXA1L-Mut, linking these changes to the copper-binding residues (Fig. 3O). YME1L depletion caused HMW species to accumulate without a proportional rise in cleavage fragments (Fig. 3P). Critically, despite stabilizing HMW OXA1L species, YME1L loss did not prevent ES-Cu-induced MRPL20 nuclear translocation (Fig. 3Q and fig. S7H) or alter ES-Cu sensitivity (fig. S7I). Thus, MRPL20 redistribution is triggered by loss of OXA1L function rather than requiring proteolytic removal of OXA1L.

### Nuclear MRPL20 promotes copper-induced ATF5 nuclear accumulation

To elucidate the nuclear function of MRPL20, we performed immunoprecipitation-mass spectrometry (IP-MS) using a nuclear-localized 2NLS-MRPL20 construct to identify its nuclear interaction partners (Fig. 4A). Quality control analyses showed the reproducibility and clear separation of control and ES-Cu samples (Fig. S8A-S8B). MRPL20 was among the most abundant proteins recovered in all samples (fig. S8C, table S6), and sensitivity analysis verified that the results were not biased by individual sample variations (fig. S8D). Copper stress markedly remodelled the MRPL20 interactome (fig. S8E, table S6), and the transcription factor ATF5 emerged as the most significantly enriched partner (Fig. 4B).

**Fig. 4.**
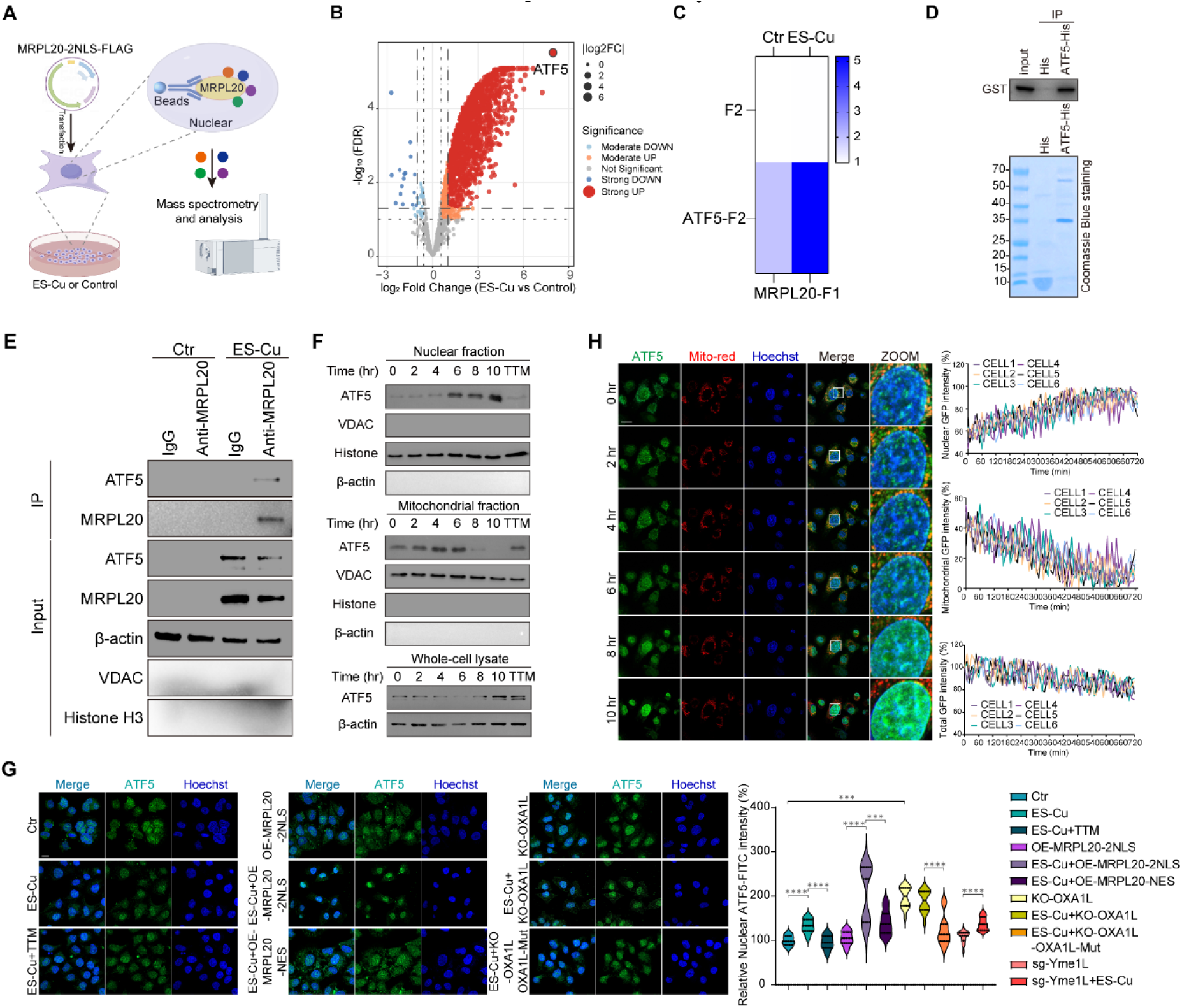
Nuclear MRPL20 binds ATF5 and promotes ATF5 nuclear accumulation during copper stress. **(A)** Schematic of immunoprecipitation–mass spectrometry (IP–MS) to identify nuclear partners of 2NLS-MRPL20–Flag in control and ES-Cu-treated cells. **(B)** Volcano plot of changes in the MRPL20 interactome after ES-Cu (1.5 μM CuSO plus 15 nM elesclomol, 10 h); ATF5 is highlighted. The axes display log fold change and −log false discovery rate. **(C)** NanoLuc binary complementation assay for the MRPL20–ATF5 interaction in HT-1080 cells under control and ES-Cu conditions. MRPL20 and ATF5 were fused to the indicated NanoLuc fragments; color represents normalized luminescence ratio (NLR). **(D)** In vitro pull-down of purified GST–MRPL20 by recombinant His-tagged ATF5; His-only protein was used as a negative control. Bound GST–MRPL20 was detected by immunoblot, and purified bait preparations are shown by Coomassie staining. **(E)** Endogenous MRPL20 co-immunoprecipitation from mitochondria-and nucleus-depleted cellular extracts under control and ES-Cu conditions. Immunoblots show co-precipitating ATF5 and the indicated input and compartment controls; IgG served as a pull-down control. **(F)** Time-course immunoblots of ATF5 in nuclear and mitochondrial fractions and whole-cell lysates following ES-Cu. TTM and VDAC, histone H3 and β-actin controls are included as indicated. **(G)** Representative ATF5 immunofluorescence images (green) and quantification of nuclear signal under the indicated ES-Cu, TTM, MRPL20 localization, OXA1L perturbation and YME1L loss conditions. Nuclei were counterstained with Hoechst (blue). Violin plots show nuclear ATF5 fluorescence from at least 10 cells per condition across three independent experiments; significance was assessed by Kruskal–Wallis test with Dunn’s correction. Scale bar, 10 μm. **(H)** Live-cell time-lapse images of ATF5–GFP (green), Mito-Red (red) and Hoechst (blue) during ES-Cu treatment, with enlarged nuclear regions at right. Traces show nuclear, mitochondrial and whole-cell ATF5–GFP fluorescence from individual cells, normalized to baseline.

We validated the physical MRPL20-ATF5 complex using complementary approaches. NanoLuc binary complementation assays detected MRPL20-ATF5 proximity in live cells and showed increased signal following ES-Cu treatment (Fig. 4C). Co-immunoprecipitation (co-IP) of tagged or endogenous proteins captured MRPL20-ATF5 complexes under untreated culture conditions (fig. S8F-S8G), and a pull-down assay using recombinant proteins supported a direct interaction between MRPL20 and ATF5 (Fig. 4D). We then asked whether copper stress promotes MRPL20-ATF5 association outside mitochondria and nuclei. Endogenous MRPL20 was immunoprecipitated from mitochondria- and nucleus-depleted extracts. An MRPL20-ATF5 complex was readily detected after ES-Cu treatment, but not in untreated controls or IgG pull-downs (Fig. 4E). These data indicate that copper stress promotes assembly of a cytoplasmic MRPL20-ATF5 complex, consistent with association before nuclear accumulation.

We next examined the temporal relationship between MRPL20 redistribution and ATF5 nuclear accumulation. ES-Cu induced a marked increase in nuclear ATF5, an effect completely blocked by the copper chelator TTM (Fig. 4F–4H). Live-cell imaging placed the onset at approximately 6 hours after ES-Cu (Fig. 4H), later than MRPL20 nuclear enrichment, which began within 4 hours (Fig. 2B and 2C). Subcellular fractionation confirmed this time-dependent rise in nuclear ATF5, with a concurrent decline in the mitochondrial fraction, whereas total ATF5 abundance increased predominantly at the later 10-hour time point (Fig. 4F). Thus, MRPL20 redistribution precedes ATF5 nuclear accumulation, which in turn precedes the later increase in total ATF5 abundance.

We then tested whether nuclear MRPL20 contributes to copper-induced ATF5 nuclear accumulation. Enforced nuclear MRPL20 (MRPL20-2NLS) did not raise nuclear ATF5 at baseline but markedly potentiated its accumulation following ES-Cu treatment. In contrast, cytoplasm-restricted MRPL20 (MRPL20-NES), which remained excluded from the nucleus during ES-Cu exposure (fig. S8H and S8I), failed to enhance nuclear ATF5 (Fig. 4G). Nuclear access of MRPL20 therefore is required for its ability to promote copper-induced ATF5 nuclear accumulation. ATF5 trafficking also tracked upstream OXA1L status. OXA1L knockout alone raised nuclear ATF5 in the absence of ES-Cu, with little further increase following copper treatment, whereas reconstituting OXA1L-knockout cells with copper-binding-deficient OXA1L-Mut restored nuclear ATF5 to near-control levels. Moreover, YME1L knockout, which stabilizes copper-damaged OXA1L without restoring its function, did not prevent ES-Cu-induced ATF5 accumulation (Fig. 4G and fig. S7J). Together, these genetic relationships place ATF5 nuclear accumulation downstream of copper-induced OXA1L inactivation and MRPL20 redistribution.

### The OXA1L-MRPL20 and ISR pathways converge on ATF5 to activate copper-induced UPR^mt^

Given ATF5’s role as a master regulator of the mitochondrial unfolded protein response (UPR^mt^) (17), we asked whether the OXA1L-MRPL20 pathway regulates UPR^mt^ transcription under copper stress. ES-Cu induced canonical UPR^mt^ transcripts, including *HSPD1*, *HSPE1*, *HSPA9*, *LONP1* and *CLPP*, in a dose-dependent manner in HeLa, HT-1080 and HepG2 cells, and this response was suppressed by the copper chelator TTM (fig. S9A). The apoptosis inducer STS or ferroptosis trigger RSL3 failed to activate the same transcripts, indicating that this signature is not a general consequence of lethal stress (fig. S9B). ATF5 knockout completely abolished this induction, whereas depletion of alternative stress transcription factors HSF1 (18) and ATF4 (19) left it intact (Fig. 5A and fig. S9C-D), establishing ATF5 as the non-redundant transcriptional driver of copper-induced UPR^mt^.

**Fig. 5.**
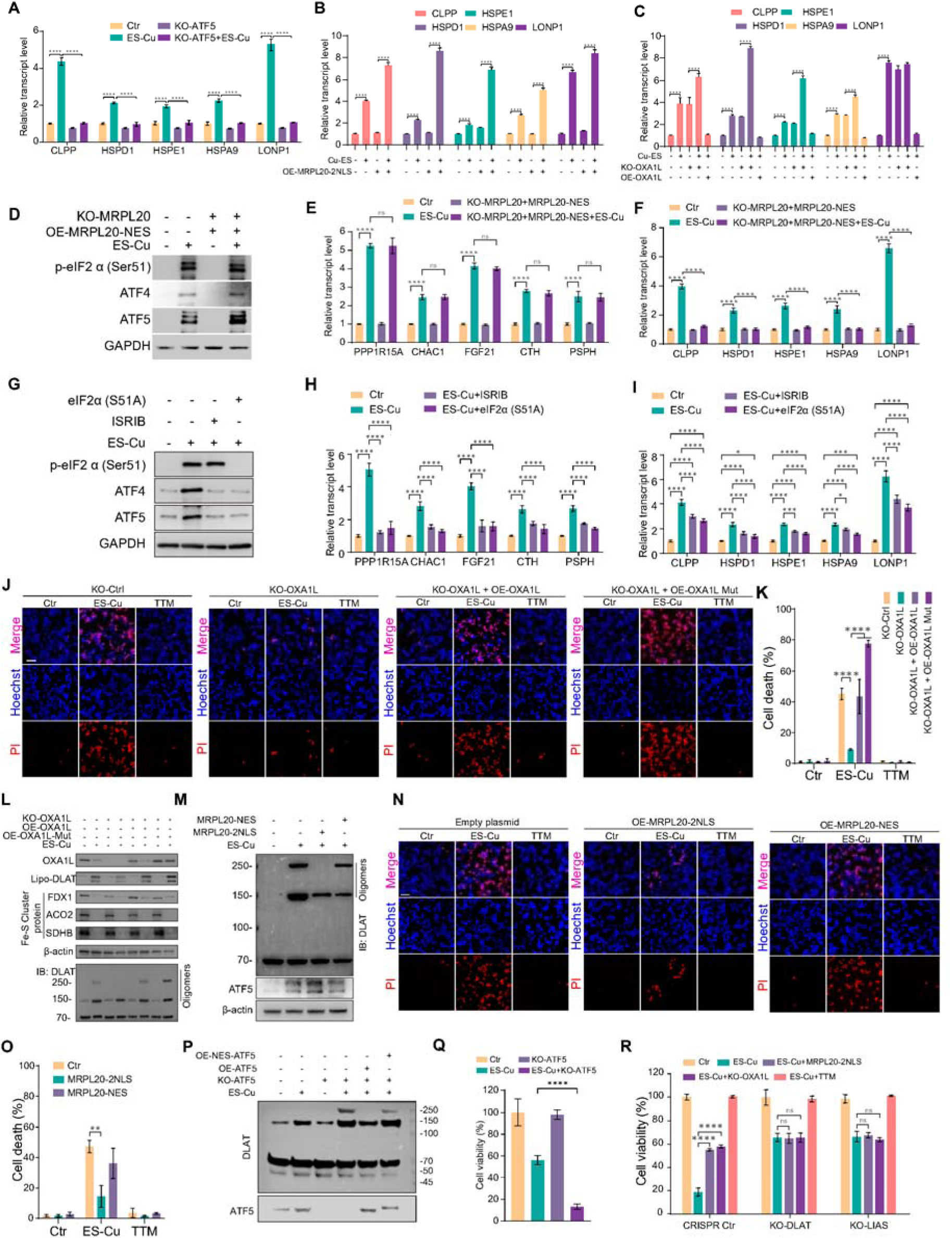
The OXA1L–MRPL20–ATF5 axis drives UPR^mt^ to restrain cuproptosis by limiting DLAT oligomer accumulation. **(A)** qPCR analysis of canonical UPR^mt^ transcripts (CLPP, HSPD1, HSPE1, HSPA9 and LONP1) in control and ATF5-knockout HT-1080 cells with or without ES-Cu. **(B)** qPCR analysis of the indicated UPR^mt^ transcripts in control and 2NLS-MRPL20-expressing HT-1080 cells with or without ES-Cu. **(C)** qPCR analysis of the indicated UPR^mt^ transcripts after OXA1L knockout or overexpression, with or without ES-Cu as shown. **(D)** Immunoblots of phosphorylated eIF2α (Ser51), ATF4 and ATF5 in MRPL20-knockout cells reconstituted with nuclear-export-signal-tagged MRPL20 (MRPL20-NES) under control and ES-Cu conditions. GAPDH is a loading control. **(E)** qPCR analysis of canonical integrated stress response (ISR) transcripts (PPP1R15A, CHAC1, FGF21, CTH and PSPH) under the MRPL20-NES conditions shown in **(D)**. **(F)** qPCR analysis of canonical UPR^mt^ transcripts under the conditions shown in **(D)**. **(G)** Immunoblots of phosphorylated eIF2α (Ser51), ATF4 and ATF5 following ES-Cu with ISRIB treatment or expression of non-phosphorylatable eIF2α(S51A). GAPDH is a loading control. **(H and I)** qPCR analysis of canonical ISR transcripts **(H)** and UPR^mt^ transcripts **(I)** under the ES-Cu, ISRIB and eIF2α(S51A) conditions shown in **(G)**. **(J)** Representative fluorescence images of Hoechst-stained nuclei (blue) and propidium iodide (PI)-positive cells (red) in control, OXA1L-knockout and OXA1L-reconstituted HT-1080 cells under control, ES-Cu and TTM conditions. Wild-type OXA1L and copper-binding-deficient OXA1L-Mut were compared. Scale bars, 50 μm. **(K)** Quantification of PI-positive cell death for the conditions shown in **(J)**. **(L)** Immunoblot analysis of OXA1L, lipoylated DLAT, the indicated Fe–S cluster proteins and DLAT oligomers after ES-Cu in control or OXA1L-perturbed cells. High-molecular-weight DLAT species were detected by anti-DLAT immunoblot under the indicated oligomer-preserving conditions. **(M)** Immunoblots of DLAT oligomeric species and total ATF5 after ES-Cu in cells expressing 2NLS-MRPL20 or MRPL20-NES; β-actin is the loading control. **(N and O)** Representative Hoechst and PI images **(N)** and quantification of PI-positive cell death **(O)** in cells expressing control vector, 2NLS-MRPL20 or MRPL20-NES under control, ES-Cu or TTM conditions. Scale bars, 50 μm. **(P)** Anti-DLAT immunoblot of oligomeric species in ATF5-knockout cells reconstituted with wild-type ATF5 or nuclear-excluded NES-ATF5, with or without ES-Cu. ATF5 expression is shown below. **(Q)** Cell viability in control and ATF5-knockout HT-1080 cells with or without ES-Cu. **(R)** CCK-8 cell viability of CRISPR control, DLAT-knockout and LIAS-knockout HT-1080 cells after ES-Cu with OXA1L knockout or enforced nuclear MRPL20, and with TTM as indicated.

We next tested whether the upstream OXA1L-MRPL20 pathway regulates this response. Enforced nuclear localization of MRPL20 was insufficient to induce UPR^mt^ transcripts at baseline, but potentiated ES-Cu-evoked transcriptional response (Fig. 5B), mirroring its effect on ATF5 nuclear accumulation. Upstream, OXA1L depletion alone raised UPR^mt^ transcripts toward ES-Cu-like levels and further enhanced the response to ES-Cu, whereas OXA1L overexpression abolished it (Fig. 5C). These findings place OXA1L and nuclear MRPL20 upstream of ATF5-dependent UPR^mt^ activation.

In parallel, copper-induced mitochondrial proteotoxic stress activates the cytosolic integrated stress response (ISR) (20), as indicated by increased eIF2α phosphorylation, ATF4 accumulation and induction of canonical ISR transcripts (fig. S9E–S9J). OXA1L depletion was sufficient to elicit a similar response, linking loss of OXA1L function to ISR activation. Because mitochondrial stress can activate the ISR through the OMA1-DELE1-HRI pathway (21), we tested the contribution of this relay. Knockout of HRI or DELE1 modestly attenuated but did not abolish, ES-Cu-induced eIF2α–ATF4 signaling and ISR-gene induction (fig. S9K and S9L), indicating that DELE1-HRI contributes to the copper-induced ISR but is not its sole upstream input.

We then asked whether MRPL20 nuclear entry is required for the ISR or specifically for UPR^mt^ activation. In MRPL20-knockout cells reconstituted with nucleus-excluded MRPL20 (MRPL20-NES), ES-Cu still induced eIF2α phosphorylation, total ATF4 and ATF5 abundance (Fig. 5D), and canonical ISR transcripts (Fig. 5E). In contrast, induction of UPR^mt^ transcripts was essentially abolished (Fig. 5F). Thus, MRPL20 nuclear localization is dispensable for activation of the ISR but is required for robust copper-induced UPR^mt^ transcription.

We next examined how the ISR contributes to ATF5 regulation. ES-Cu raised ATF5 abundance through two established ISR-responsive mechanisms: CHOP-dependent transcription of *ATF5* (22) (fig. S9M and S9N) and eIF2α-dependent translational control through upstream open reading frames in the *ATF5* 5′ leader (23) (fig. S9O–S9R). Consistent with these mechanisms, blocking ISR signalling with ISRIB (24) or non-phosphorylatable eIF2α(S51A) attenuated the increase in ATF5 abundance and reduced, but did not abolish, UPR^mt^ transcripts (Fig. 5G–5I and fig. S9S). Moreover, ISRIB did not further suppress UPR^mt^ transcripts in ATF5-deficient cells (fig. S9S), consistent with ATF5 mediating the ISR-dependent component of UPR^mt^ activation.

Together, these data define two mechanistically separable pathways that converge on ATF5. The ISR increases ATF5 abundance through transcriptional and translational regulation, whereas the OXA1L-MRPL20 pathway promotes its effective nuclear accumulation and is specifically required for UPR^mt^ transcriptional output. Their convergence enables robust UPR^mt^ activation during copper stress while the ISR cannot substitute for the OXA1L-MRPL20 branch.

### UPR^mt^ restrains DLAT oligomerization and cuproptosis

We next asked how UPR^mt^ activation affects cuproptosis. OXA1L knockout, which strongly activates UPR^mt^, significantly reduced ES-Cu-induced cell death. Re-expression of wild-type OXA1L restored ES-Cu cytotoxicity, whereas reconstitution with the copper-binding-deficient OXA1L-Mut, which blunts UPR^mt^ activation, exacerbated death relative to controls (Fig. 5J and 5K and fig. S10A). These genetic perturbations support UPR^mt^ activation as an endogenous pro-survival mechanism to counter cuproptosis. Examining canonical cuproptosis hallmark events, we found that increased DLAT lipoylation and reduction of Fe-S cluster proteins (FDX1, ACO2, SDHB) were unaffected by OXA1L status. Strikingly, OXA1L knockout markedly suppressed the accumulation of high-molecular-weight DLAT oligomers. Conversely, reconstitution with the UPR^mt^-blunting OXA1L-Mut enhanced DLAT oligomerization (Fig. 5L). Thus, activation of the OXA1L-dependent UPR^mt^ does not reverse the upstream copper-induced metabolic alterations but instead limits accumulation of oligomerized DLAT, a major proteotoxic feature of cuproptosis.

Perturbation of the downstream MRPL20-ATF5 axis produced concordant effects. Enforced nuclear localization of MRPL20 (MRPL20-2NLS) efficiently inhibited ES-Cu-induced DLAT oligomerization without an evident change in total ATF5 across the MRPL20 localization conditions (Fig. 5M), and reduced cell death (Fig. 5N and 5O and fig. S10B), whereas nucleus-excluded MRPL20 (MRPL20-NES) did not confer comparable protection (Fig. 5N and 5O). ATF5 knockout substantially increased DLAT oligomer accumulation following ES-Cu treatment, and reconstitution with wild-type ATF5 suppressed this phenotype. By contrast, expression of nuclear-excluded ATF5 (NES-ATF5) failed to restore protection (Fig. 5P), showing that protection requires nuclear ATF5. ATF5 knockout also increased sensitivity to ES-Cu, as shown by a further reduction in cell viability compared with control cells (Fig. 5Q). Together, these results link nuclear MRPL20 and ATF5-dependent UPR^mt^ activity to suppression of copper-induced DLAT oligomer accumulation and cell death.

To further determine whether the protective effect of OXA1L-MRPL20 pathway depends on the canonical lipoylated-protein branch of cuproptosis, we examined DLAT- and LIAS-deficient cells. Neither OXA1L knockout nor enforced nuclear MRPL20 provided additional protection from ES-Cu in these backgrounds (Fig. 5R), consistent with this pathway acting within the lipoylated DLAT-dependent branch of cuproptosis. As an independent test, the electron transport chain inhibitors rotenone and antimycin A induced UPR^mt^ transcripts and were accompanied by reduced DLAT oligomerization and decreased ES-Cu-induced cytotoxicity (fig. S11). Because these agents broadly alter mitochondrial metabolism, these findings provide supportive rather than definitive evidence for UPR^mt^-mediated protection. Together with the genetic perturbation of OXA1L-MRPL20-ATF5 pathway, these data support a model in which UPR^mt^ activation limits copper-induced DLAT oligomer accumulation and raises the threshold for cuproptotic cell death.

### Inhibition of UPR^mt^ sensitizes tumors to copper-induced cytotoxicity

To assess the therapeutic relevance of the OXA1L-MRPL20-ATF5-UPR^mt^ pathway, we first examined the expression of its components across human malignancies using the GEPIA3 database (25). MRPL20, ATF5 and several downstream UPR^mt^ effectors, including the chaperones HSPD1, HSPA9, DNAJA3 and HSPE1 and the protease LONP1, show increased transcript abundance in multiple tumor types relative to corresponding normal tissues, including colon adenocarcinoma (fig. S12A and S12B, table S7). Although these expression patterns do not establish pathway activity, they provided a rationale to test whether pharmacological inhibition of the UPR^mt^ could enhance sensitivity to copper stress.

DCEM1 has been reported to inhibit UPR^mt^ by directly binding HSP60 and disrupting its interaction with the protease CLPP (26). At 10 µM, DCEM1 alone had little effect on cell viability (fig. S12C) but reduced CLPP protein abundance, as did HSP60 knockout (Fig. 6A). Neither DCEM1 treatment nor HSP60 knockout evidently reduced total ATF5 abundance (Fig. 6A). To localize its action, we examined the two trafficking events upstream of UPR^mt^. DCEM1 reduced ES-Cu-induced nuclear ATF5 accumulation (Fig. 6B) without preventing MRPL20 nuclear translocation (Fig. 6C and fig. S12E). Consistent with this placement, DCEM1 suppressed induction of canonical UPR^mt^ transcripts during ES-Cu treatment (Fig. 6D) and increased accumulation of high-molecular-weight DLAT oligomers (Fig. 6E and fig. S12D). DCEM1 therefore acts downstream of MRPL20 retrograde trafficking, at the level of nuclear ATF5.

**Fig. 6.**
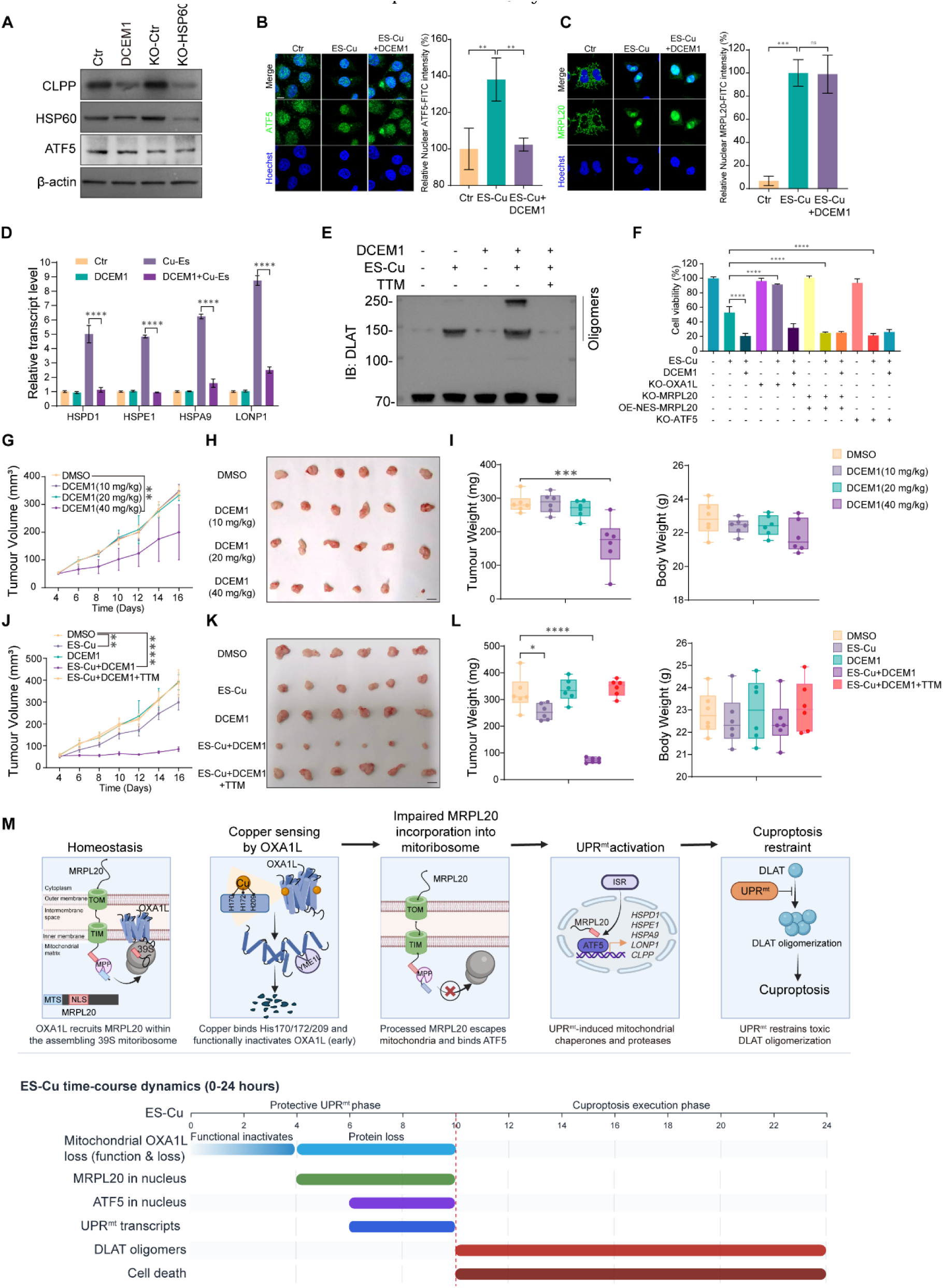
UPR^mt^ inhibition sensitizes tumors to copper-induced cytotoxicity. **(A)** Immunoblots of CLPP, HSP60 and total ATF5 in control, DCEM1-treated, knockout-control and HSP60-knockout cells; β-actin is the loading control. **(B and C)** Representative immunofluorescence images and quantification of nuclear endogenous ATF5 **(B)** or MRPL20 **(C)** in control, ES-Cu and ES-Cu plus DCEM1 conditions; Hoechst marks nuclei. **(D)** qPCR analysis of HSPD1, HSPE1, HSPA9 and LONP1 transcripts in HCT-116 cells treated with ES-Cu (1.5 μM CuSO plus 15 nM elesclomol), DCEM1 (10 μM) or their combination. **(E)** Anti-DLAT immunoblot of high-molecular-weight oligomeric species in cells treated with the indicated combinations of ES-Cu, DCEM1 and TTM. **(F)** Cell viability of HCT-116 cells following ES-Cu and DCEM1 treatment under the indicated OXA1L-knockout, MRPL20-knockout with MRPL20-NES re-expression, or ATF5-knockout conditions. **(G)** Growth curves of subcutaneous HCT-116 xenografts following treatment with vehicle (DMSO) or DCEM1 at 10, 20 or 40 mg/kg. n = 6 mice per group. **(H)** Endpoint photographs of tumors from the dose-response study in **(G)**. **(I)** Endpoint tumor weights (left) and mouse body weights (right) for the groups in **(G)**. **(J)** Tumor growth curves in HCT-116 xenografts given vehicle, ES-Cu [ES (20 mg/kg) plus CuSO (0.5 mg/kg)], DCEM1 (20 mg/kg), ES-Cu plus DCEM1, or ES-Cu plus DCEM1 and TTM (10 mg/kg). n = 6 mice per group. **(K)** Endpoint photographs of tumors from the combination-treatment study in **(J)**. **(L)** Endpoint tumor weights (left) and mouse body weights (right) for the groups in **(J)**. **(M)** Proposed working model and approximate time course after ES-Cu. Copper binding functionally inactivates OXA1L, MRPL20 is released from the assembling 39S mitoribosomal subunit and a processed MRPL20 pool appears in the nucleus, where it associates with ATF5 and supports a protective UPR^mt^ program that limits DLAT oligomer accumulation. The integrated stress response (ISR) also contributes to ATF5 regulation. The route by which MRPL20 leaves mitochondria remains unresolved.

DCEM1 enhanced ES-Cu cytotoxicity across HepG2, LNCaP, A549 and SUM159PT cells with TTM restoring viability in each model, indicating that this sensitization was copper dependent (fig. S12F-S12I). In HCT-116 cells, combined DCEM1 and ES-Cu treatment reduced viability and increased cell death more strongly than either treatment alone, whereas TTM reversed this effect (fig. S12J-S12L).

To test whether this requires the OXA1L-MRPL20-ATF5 axis, we disrupted the pathway genetically in HCT-116 cells (Fig. 6F). None of the perturbations altered baseline viability. ES-Cu alone reduced viability to 53%, and OXA1L knockout conferred near-complete protection while replacement of endogenous MRPL20 with nuclear-excluded NES-MRPL20 reduced viability to 25%, and ATF5 knockout to 21%. Adding DCEM1 to ES-Cu-treated control cells reduced viability to 20%, phenocopying genetic blockade at either downstream node. DCEM1 abolished the protection conferred by OXA1L loss but produced no further sensitization in NES-MRPL20 cells or ATF5-knockout cells. DCEM1 therefore acts downstream of OXA1L and requires the MRPL20-ATF5 arm for its sensitizing activity.

We next tested this strategy in HCT-116 xenografts. DCEM1 alone had little effect on tumor growth at 10 or 20 mg/kg, whereas 40 mg/kg reduced tumor growth; body weight changed minimally across these doses (Fig. 6G-6I). We therefore evaluated DCEM1 at 20 mg/kg in combination with ES (20 mg/kg) and CuSO (0.5 mg/kg) (fig. S12M). The combination suppressed tumor growth and reduced endpoint tumor weight more strongly than either treatment alone, and addition of TTM reversed this effect, supporting copper dependence (Fig. 6J–6L). No substantial differences in body weight were detected among these treatment groups (Fig. 6L). Together, these findings show that pharmacological inhibition of the UPR^mt^ removes a protective constraint on copper-induced DLAT oligomer accumulation and sensitizes tumors to copper-induced cytotoxicity.

## Discussion

Mitochondrial proteotoxic stress is a defining feature of cuproptosis (3), yet how cells sense copper-evoked mitochondrial proteotoxicity and mount a defensive response remains poorly understood. Here, we define a retrograde signaling cascade that couples copper sensing at the inner mitochondrial membrane to a nuclear UPR^mt^ transcriptional program (Fig. 6M). OXA1L is best known as an evolutionarily conserved insertase that coordinates mitochondrial translation with co-translational membrane insertion and respiratory-chain assembly (8, 27, 28). Our findings reveal a previously unrecognized moonlighting function of OXA1L in copper-stress perception. Copper binding requires His170, His172 and His209 and disrupts OXA1L homodimerization and its association with the large mitoribosomal subunit. Importantly, preventing OXA1L degradation through YME1L loss does not block downstream MRPL20 redistribution, indicating that loss of OXA1L function, rather than its proteolytic clearance, initiates signaling. How copper binding alters OXA1L structure remains unresolved and will require direct structural analysis. Nevertheless, these findings position OXA1L as a copper-responsive mitochondrial sensor that links metal overload to disruption of mitochondrial protein homeostasis.

MRPL20 emerges from this response as a non-canonical stress messenger that relays mitochondrial dysfunction to the nucleus. This process differs from passive precursor-overflow models in which generalized import failure causes accumulation of unprocessed mitochondrial proteins in the cytosol (18, 29, 30). Three observations argue against such a mechanism. CHX strongly suppressed MRPL20 nuclear accumulation, indicating that the nuclear pool depends on ongoing protein synthesis, whereas dual-end tagging showed loss of the N-terminal tag with retention of the C-terminal tag in the nuclear MRPL20, consistent with mitochondrial import and presequence processing before its nuclear appearance (13). Copper-induced release of MRPL20 from isolated mitochondria and loss of its association with the 39S component MRPL21, both suppressed by copper-binding-deficient OXA1L-Mut, further support a model in which OXA1L inactivation impairs stable incorporation of MRPL20 into the assembling 39S subunit (16), making a processed MRPL20 pool available for release. The absence of nuclear accumulation of MRPL48 and MRPL51, together with continued import of a presequence-dependent mitochondrial reporter, argues against generalized mitochondrial protein escape or global collapse of mitochondrial import. MRPL20’s intrinsic NLS then directs the exported pool to the nucleus.

The route by which processed MRPL20 traverses the mitochondrial membranes remains unresolved. Nevertheless, regulated protein exit from intact mitochondria has clear precedent. Processed fumarase and PINK1 both reappear in the cytosol after mitochondrial import and proteolytic maturation (31, 32), and sulfite oxidase maturation illustrates how presequence-dependent import, proteolytic processing, and folding-based retention can be mechanistically coupled (33). Protein export also contributes to mitochondrial proteostasis. Conformationally destabilized proteins in the intermembrane space can be released through TOM40 (34), and matrix quality-control substrates were shown in a cell-free system to undergo ATP-dependent retrotranslocation across both mitochondrial membranes with TOM40 contributing to their passage across the outer membrane (35). Stress-induced mitochondria-to-nucleus translocation has also been reported for GPS2 (36) and selected TCA-cycle dehydrogenases (37). These findings establish that selective mitochondrial protein export is biologically plausible, but they do not imply that MRPL20 uses the same route. Defining the machinery that mediates its release and couples mitochondrial exit to nuclear import will be important for understanding whether MRPL20 represents a dedicated pathway or one example of a broader class of mitochondrial retrograde messengers.

Our findings also refine the current model of mammalian UPR^mt^ signaling. By analogy with *C. elegans* ATFS-1, stress-induced mitochondrial import failure is thought to redirect ATF5 to the nucleus (17). Our data are consistent with such a shift in ATF5 partitioning as mitochondrial ATF5 declined while its nuclear abundance rose. How ATF5 translates diverse mitochondrial stresses into a defined UPR^mt^ output, however, has remained unclear. We find that this output cannot be explained by ATF5 availability or nuclear localization alone. When endogenous MRPL20 is replaced with a nuclear-excluded form (MRPL20-NES), total ATF5 induction and ATF5 nuclear accumulation all remain intact, yet UPR^mt^ transcription is strongly attenuated. Conversely, nuclear-targeted MRPL20 (MRPL20-2NLS) potentiates both nuclear ATF5 and UPR^mt^ output under copper without raising total ATF5. Nuclear MRPL20 may therefore provide a tuning mechanism for this response, acting alongside with ATF5 as a transcriptional co-activator, a possibility require further investigation. These findings extend an emerging principle in which mitochondrial ribosomal proteins can acquire signaling functions outside their canonical role in ribosome biogenesis (38, 39) and they provide a mechanism linking a defined mitochondrial perturbation to mammalian nuclear stress signaling.

A further conceptual advance is the interaction of this OXA1L-MRPL20 pathway with the ISR during copper stress. Rather than forming a simple linear hierarchy, the data support two mechanistically separable inputs that converge on ATF5. The OXA1L-MRPL20 pathway promotes effective ATF5 nuclear accumulation and is specifically required for robust UPR^mt^ transcriptional output, whereas the ISR reinforces output through transcriptional and translational control of ATF5 abundance. Accordingly, broad ISR blockade only partially reduces UPR^mt^ output, whereas preventing MRPL20 nuclear entry or deleting ATF5 abolishes it. Convergence of these pathways on ATF5 therefore provides a mechanism by which mitochondrial copper stress can couple a specific retrograde signal to broader stress-dependent amplification of the UPR^mt^.

Functionally, this pathway constitutes an endogenous defense against cuproptosis. It does not antagonize copper-mediated lipoylation or loss of Fe-S proteins. Instead, it mitigates the execution phase by restraining the accumulation of high-molecular-weight DLAT oligomers, the proteotoxic hallmark of cuproptosis (3). A recent study identified copper-induced polymerization of MYH9 as an additional execution mechanism for cuproptosis and showed that DLAT-dependent acetylation facilitates MYH9 copper binding (40). However, whether this execution pathway interacts with UPR^mt^-dependent control of DLAT oligomerization remains unknown. How the UPR^mt^ restrains DLAT oligomers also remains unknown. Determining which ATF5-induced chaperones or proteases directly limit DLAT oligomer accumulation should therefore help define how mitochondrial proteostasis sets the threshold for cuproptotic death.

The protective nature of this pathway also creates a potential therapeutic vulnerability. Pharmacological inhibition of UPR^mt^ with DCEM1 enhanced ES-Cu-induced DLAT oligomerization and cell death across multiple tumor cell models and increased the antitumor effect of ES-Cu in HCT-116 xenografts. These findings provide proof of principle that disabling the mitochondrial proteostasis response can increase sensitivity to copper overload. However, DCEM1 acts on the HSP60-CLPP proteostasis machinery rather than specifically on the OXA1L-MRPL20 pathway, and the in vivo combination has so far been evaluated in a single xenograft model. Establishing the generality and therapeutic window of this strategy will therefore require additional tumor models and orthogonal genetic or pharmacological approaches to UPR^mt^ inhibition.

Collectively, our work uncovers OXA1L as a mitochondrial copper sensor and defines a mitochondria to nucleus retrograde signaling cascade that shapes cellular susceptibility toward cuproptosis, offering new perspectives for targeting copper driven proteotoxicity in tumors.

## Supporting information

All Table

## Acknowledgments

This research was funded by National Natural Science Foundation of China (No. 82473894) to C, F; National Natural Science Foundation of China (No.82073412) to Q, Z; J. K received support from the 5Point Foundation (Christine Martin Fellowship).

## Author contributions

Conceptualization: CF, QZ and JK. Methodology: CF, QZ and XY.

Investigation: XY, CW, GS, SZ, KZ and XD. Formal analysis: CS and XL.

Visualization: XY and CW.

Funding acquisition: CF, QZ and JK. Supervision: CF.

Writing – original draft: XY.

Writing – review & editing: CF, JK, XY, CS.

## Competing interests

Authors declare that they have no competing interests.

## Data and materials availability

All data are available in the main text or the supplementary materials.

## Supplementary Materials

Materials and Methods Figs. S1 to S12

Tables S1 to S8

## Notes

### Competing Interest Statement

The authors have declared no competing interest.

