## SupplementalMaterial for "Copper sensing by OXA1L triggers a mitochondria-to-nucleus stress response to restrain cuproptosis"

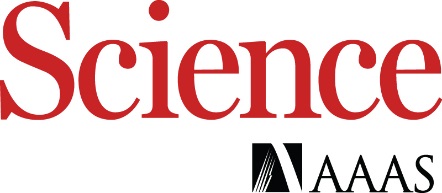

Supplementary Materials for

**Copper sensing by OXA1L triggers a mitochondria-to-nucleus stress response to restrain cuproptosis**

Xueliang Yang^1,2^†, Chong Wei^1^†, Ye Lu^3^†, Qinglin Li^4^†, Xin Liu^3^, Gang Shao^1^, Shuyuan Zhang^5^, Hengzhi Li^1^, Yuzhe Wang^1^, Keke Zheng^1^, Xiaoyan Dai^1^, Jian Kang^6,7^*, Chulin Sha^3^*, Qiang Zhang^2^*, Caiyun Fu^1#^*

**The PDF file includes:**

Materials and Methods

Figs. S1 to S12

**Other Supplementary Materials for this manuscript include the following:**

Tables S1 to S8

Materials and Methods

Cell lines and cell culture

A panel of human cancer cell lines, including HT-1080 (fibrosarcoma), A549 (lung carcinoma), HeLa (cervical carcinoma), HepG2 (hepatocellular carcinoma), and U2OS (osteosarcoma), were commercially sourced from the Cell Bank of the Chinese Academy of Sciences. The HEK293T (human embryonic kidney) cell line was a generous gift from Dr. Ge Bai (Zhejiang University School of Medicine). Cells were cultured in high-glucose DMEM (Meilunbio) supplemented with 10% heat-inactivated FBS (ExCell Bio) and 1% penicillin-streptomycin (Pricella). Cultures were kept in a Thermo Scientific Forma series incubator at 37°C with a 5% CO₂ atmosphere and passaged at 80-90% confluence using trypsin-EDTA. All cell lines were authenticated and consistently tested negative for mycoplasma.

Isolation and Culture of Mouse Articular Chondrocytes

Animal experiments were approved by the Institutional Animal Care and Use Committee (IACUC) of Zhejiang university (Approval No. ZJU20250707). Mice were housed under standard conditions (12 h light/dark cycle, ad libitum access to food and water).

Primary murine articular chondrocytes were isolated as previously described. Briefly, bilateral hip joints were harvested from 4-week-old C57BL/6 mice (weighing 14.0 ± 2.0 g) under sterile conditions. Articular cartilage was dissected, minced into approximately 1 mm³ pieces, and digested sequentially. First, the tissue fragments were treated with 0.25% trypsin for 30 minutes at 37°C. Subsequently, they were digested with 0.2% (w/v) type II collagenase (Sigma-Aldrich, USA) on a horizontal shaker at 37°C for 4 hours. The resulting cell suspension was filtered through a 70-μm cell strainer to remove debris and then centrifuged at 300 × g for 5 minutes. The pelleted chondrocytes were resuspended and seeded into culture flasks containing Dulbecco‘s Modified Eagle Medium supplemented with 10% fetal bovine serum and 1% penicillin-streptomycin. Cells were maintained at 37°C in a humidified atmosphere containing 5% CO₂. The initially attached cells were designated as passage 0 (P0). The chondrocyte phenotype was identified by their characteristic polygonal, cobblestone-like morphology.

Isolation and Culture of Mouse Embryonic Fibroblasts (MEFs)

Primary mouse embryonic fibroblasts (MEFs) were isolated from ICR mouse embryos at embryonic day 13.5 (E13.5). Briefly, pregnant mice were euthanized by cervical dislocation under protocols approved by the Institutional Animal Care and Use Committee (IACUC) of Zhejiang university (Approval No. ZJU20250707). The uterine horns were aseptically dissected and transferred to a sterile dish containing cold phosphate-buffered saline (PBS). Individual embryos were separated from the embryonic membranes and placenta. After removal of the head, limbs, tail, and visceral tissues (heart, liver, intestines), the remaining embryonic torsos were thoroughly rinsed with PBS. The tissues were then minced into fine pieces (approximately 1 mm³) using sterile surgical blades. The tissue fragments were digested with 0.25% trypsin-EDTA (Gibco, USA) at 37°C for 20 minutes with gentle agitation. Digestion was terminated by adding an equal volume of complete growth medium: Dulbecco’s Modified Eagle Medium (DMEM, Gibco, USA) supplemented with 10% fetal bovine serum (FBS, Gibco, USA) and 1% penicillin-streptomycin (Gibco, USA). The cell suspension was filtered through a 70-μm cell strainer to remove undigested tissue clumps and then centrifuged at 300 × g for 5 minutes. The cell pellet was resuspended in complete growth medium and seeded into culture-treated flasks. Cells from this initial plating were designated as passage 0 (P0). MEFs were cultured at 37°C in a humidified atmosphere with 5% CO₂ and passaged at approximately 80-90% confluence. Early-passage MEFs (passages 1-3, P1-P3) were used for subsequent experiments.

Plasmid Construction

All oligonucleotide sequences used in this study (see Table S8 for complete list) were synthesized by Tsingke Biotechnology Co., Ltd. (Beijing, China). The backbone vectors, including pcDNA3.1(+), pCDH-CMV-EF1α, pLKO.1-TRC, lentiCRISPR v2, and pET-28a (+), were obtained from Addgene. All gene fragments for cloning were amplified from a cDNA library derived from HT-1080 human fibrosarcoma cells.

To construct plasmids for gene overexpression, the full-length coding sequences (CDS) of the target genes were amplified via high-fidelity PCR using Phusion polymerase (New England Biolabs) with gene-specific primers flanked by appropriate restriction sites. The PCR amplicons and the recipient vectors (pcDNA3.1(+) or pCDH-CMV-EF1α-EGFP) were digested with NheI and EcoRI. The digested fragments were purified and ligated using the ClonExpress Ultra One Step Cloning Kit (C112, Vazyme) according to the manufacturer's protocol, and the ligation products were transformed into competent E. coli DH5α cells for amplification and plasmid isolation.

For recombinant protein expression and purification, the CDS of ATF5, DNAJA3, HSPA9, HSPD1, and HSPE1 were separately cloned into the pET-28a (+) vector. Each gene fragment was amplified with primers designed to fuse it in-frame with an N-terminal 6×His tag encoded by the vector, resulting in constructs (6×His-ATF5, -DNAJA3, -HSPA9, -HSPD1, and -HSPE1) suitable for isopropyl β-D-1-thiogalactopyranoside (IPTG)-inducible expression in E. coli (BL21 DE3 chemically competent cell, Tsingke, TSC-E01).

For RNA interference, shRNA knockdown plasmids were generated by annealing complementary single-stranded oligonucleotides to create double-stranded DNA fragments. These fragments were then ligated into the AgeI and EcoRI-linearized pLKO.1 vector using T4 DNA ligase (C301, Vazyme), producing constructs that express shRNAs under the control of the U6 promoter.

For CRISPR/Cas9-mediated gene knockout, single-guide RNA (sgRNA) sequences targeting specific genomic loci were designed. The annealed sgRNA oligonucleotides were cloned into the BsmBI-v2-digested lentiCRISPR v2 backbone using T4 DNA ligase, resulting in a vector that co-expresses the sgRNA and the Cas9 nuclease.

To generate organelle-specific proximity labeling constructs for subsequent applications, DNA sequences encoding the V5 epitope tag and the miniTurboID enzyme were fused in-frame to the N- or C-terminus of various organelle-targeting bait proteins or signal peptides. These fusion cassettes were subsequently cloned into the pCDH-CMV-EF1α lentiviral vector for stable expression in mammalian cells.

All constructed plasmids were verified by restriction enzyme digestion and Sanger sequencing performed by Tsingke Biotechnology Co., Ltd. Plasmid DNA for transfection was prepared using the HiSpeed Plasmid Maxi Kit (Qiagen) according to the manufacturer's instructions, and its concentration and purity were assessed by spectrophotometry.

Lentivirus Production and Stable Cell Line Generation

Lentiviruses were packaged in HEK-293T cells. Briefly, HEK-293T cells were seeded in 60-mm culture dishes at a density of 1 × 10⁶ cells/dish in complete growth medium and incubated overnight to reach 70-90% confluence at the time of transfection. For viral packaging, the cells were co-transfected with a 4:3:1 mass ratio of the lentiviral transfer vector (either pCDH-CMV-MCS-EF1-Puro or pLKO.1-Puro), the packaging plasmid (psPAX2), and the envelope plasmid (pMD2.G) using the Liposomal Transfection Reagent (Cat# 40802ES02, Yeasen) according to the manufacturer’s protocol. The transfection complex was prepared in Opti-MEM medium and added dropwise to the cells.

48 to 72 hours post-transfection, the virus-containing supernatant was collected, passed through a 0.45-μm pore-size sterile filter to remove cellular debris, and either used immediately or aliquoted and stored at -80°C. For transduction, target cells (e.g., HT-1080) were seeded in 6-well plates at ~30% confluence and infected with the filtered viral supernatant in the presence of polybrene (Yeasen) at a final concentration of 10 μg/mL. To achieve optimal infection efficiency, the multiplicity of infection (MOI) was empirically determined for each cell line and viral batch. After 24 hours of incubation, the viral supernatant was replaced with fresh complete medium.

48 hours post-transduction, the cells were subjected to antibiotic selection to establish stable polyclonal populations. Cells transduced with pCDH-based vectors were selected with 2-4 μg/mL puromycin (Invitrogen), while cells transduced with pLKO.1-based shRNA vectors were selected with 1-2 μg/mL puromycin. The selection medium was refreshed every 2-3 days, and the process was continued for 1 week until all non-transduced control cells were completely dead. The surviving cells were considered as a stable polyclonal pool.

For the key mechanistic experiments, knockdown or overexpression was assessed by western blotting or quantitative RT-PCR (qRT-PCR), as shown in the corresponding figures. The discovery screen was used to prioritize candidates and did not include individual protein-level validation of every shRNA or overexpression construct. All shRNA sequences used in this study are listed in Table S8. The stable cells were maintained under appropriate antibiotic pressure and used for subsequent experiments within 10 passages to minimize phenotypic drift.

CRISPR/Cas9-Mediated Gene Knockout

Single-guide RNA (sgRNA) sequences targeting the human OXA1L gene and other genes of interest were designed using the online CHOPCHOP tool (http://chopchop.cbu.uib.no). For each target, two independent sgRNA sequences were selected to mitigate off-target effects. The oligos encoding these sgRNAs (sequences listed in Table S8) were annealed and cloned into the BsmBI-v2 restriction site of the lentiCRISPR v2 vector. All constructs were verified by Sanger sequencing. To produce lentivirus, the sgRNA-expressing lentiCRISPR v2 vector was co-transfected with the packaging plasmid psPAX2 and the envelope plasmid pMD2.G into HEK-293T cells using Liposomal Transfection Reagent (Yeasen). Viral supernatants were collected 48-72 hours post-transfection, filtered through a 0.45-μm filter, and used to infect target cells (e.g., HT-1080) in the presence of polybrene (8 μg/mL). At 48 hours post-infection, cells were selected with puromycin (2 μg/mL) for 5-7 days to obtain a polyclonal population of transduced cells.

For the generation of knockout cell lines, such as for OXA1L, a single-cell cloning step was performed. The puromycin-selected polyclonal population was trypsinized, serially diluted, and seeded into 96-well plates at a density of approximately 0.5 cells per well to isolate single-cell-derived clones. Individual clones were expanded over 2-3 weeks. It is noteworthy that OXA1L knockout led to a significant reduction in cellular proliferation. Therefore, after initial expansion, the positive clones were maintained and passaged for an additional week to allow the cell population to stabilize and regain a consistent growth rate before being used in any subsequent functional assays. Knockout efficiency was confirmed at the protein level by Western blotting and at the genomic level by Sanger sequencing of the target locus following PCR amplification.

For all other target genes besides OXA1L, a pooled knockout strategy was employed using a polyclonal cell population. To enhance knockout efficiency and compensate for potential variation in individual sgRNA activity, cells were transduced with a lentiviral pool containing both independent sgRNAs targeting the same gene. Following puromycin selection, the entire polyclonal cell pool was used directly for subsequent experiments. The knockout efficiency in these pooled populations was confirmed by Western blotting to demonstrate a substantial reduction in target protein expression. All established knockout cell lines and pools were maintained under continuous puromycin selection and used within 10 passages to ensure phenotypic consistency.

Antibodies

The following primary antibodies were used for immunoblotting (IB), immunohistochemistry (IHC), and immunofluorescence (IF): anti-OXA1L (Proteintech, 21055-1-AP, RRID: AB_10695769; IB 1:1000, IHC 1:200), anti-MRPL20 (Affinity Biosciences, DF3663, RRID: AB_2836035; IB 1:1000, IF 1:300-500), anti-DLAT (Cell Signaling Technology, 12362T, RRID: AB_2797893; IB 1:1000), anti-FDX1 (Proteintech, 12592-1-AP, RRID: AB_11182486; IB 1:1000, IHC 1:500), anti-ACO2 (Proteintech, 11134-1-AP, RRID: AB_2289288; IB 1:1000), anti-SDHB (Proteintech, 67600-1-Ig, RRID: AB_2882806; IB 1:1000), anti-HSP60 (Proteintech, 15282-1-AP, RRID: AB_2121440; IB 1:1000), anti-ClpP (Proteintech, 15698-1-AP, RRID: AB_2245115; IB 1:1000, IHC 1:200), anti-ATF5 (Proteintech, 67066-1-Ig, RRID: AB_2882376; IB 1:1000, IF 1:300), anti-VDAC (Proteintech, 81538-1-RR, RRID: AB_2935560; IB 1:1000), anti-Histone H3 (Proteintech, 17168-1-AP, RRID: AB_2716755; IB 1:1000), anti-HSP70 (Proteintech, 10995-1-AP, RRID: AB_2264230; IB 1:1000), anti-LONP1 (Proteintech, 15440-1-AP, RRID: AB_2137152; IB 1:1000), anti-OMA1 (Proteintech, 17116-1-AP, RRID: AB_2299053; IB 1:1000), anti-Yme1L (Proteintech, 11510-1-AP, RRID: AB_2217459; IB 1:1000), anti-AFG3L2 (Proteintech, 14631-1-AP, RRID: AB_2242420; IB 1:1000), anti-Lipoic acid (Abcam, ab58724, RRID: AB_880635; IB 1:1000), anti-Streptavidin (Cell Signaling Technology, 3999S, RRID: AB_10830897; IB 1:5000), anti-DYKDDDDK tag (Proteintech, 66008-4-Ig, RRID: AB_2918475; IB 1:5000), anti-V5 tag (Proteintech, 14440-1-AP, RRID: AB_2878059; IB 1:5000), anti-HA tag (Proteintech, 51064-2-AP, RRID: AB_11042321; IB 1:5000), anti-His tag (Proteintech, 66005-1-Ig, RRID: AB_11232599; IB 1:5000), anti-GST tag (Proteintech, 10000-0-AP, RRID: AB_11042316; IB 1:5000), anti-β-Actin (ABclonal, AC026, RRID: AB_2768234; IB 1:10000), and anti-GAPDH (ABclonal, A19056, RRID: AB_2768234; IB 1:10000). HRP-conjugated goat anti-mouse IgG (H+L) (ABclonal, AS003, RRID: AB_2769851; IB 1:10000) and HRP-conjugated goat anti-rabbit IgG (H+L) (ABclonal, AS014, RRID: AB_2769854; IB 1:10000) were used as secondary antibodies for immunoblotting. For immunofluorescence, Alexa Fluor 488-conjugated anti-rabbit IgG (H+L) (Proteintech, SA00013-2, RRID: AB_2797132; IF 1:500) and FITC-conjugated anti-mouse IgG (H+L) (Proteintech, SA00003-2, RRID: AB_2890897; IF 1:500) were applied.

Reagents and Compounds

Elesclomol (HY-12040), biotin (HY-B0511), dithiothreitol (DTT, HY-15917), DCEM1 (HY-171616), rotenone (HY-B1756), staurosporine (STS, HY-15141), (1S,3R)-RSL3 (HY-100218A), erastin (HY-15763), MG132 (HY-13259), chloroquine (CQ, HY-17589A), Z-VAD-FMK (HY-16658B), and MDL-28170 (HY-18236) were purchased from MedChemExpress (MCE). Copper(II) sulfate pentahydrate (CuSO₄·5H₂O) (451657), antimycin A (A8674), and isopropyl β-D-1-thiogalactopyranoside (IPTG, I6758) were obtained from Sigma-Aldrich. DTSP (57757-57-0) was sourced from Solarbio. Puromycin (ant-pr-1) was purchased from InvivoGen, and blasticidin S (ST018) was from Beyotime. Hoechst 33342 (C1022) was obtained from Beyotime, and propidium iodide (PI, 40710ES03) was from Yeasen. Furimazine (E5806) was purchased from Selleck.

Materials.

Total RNA was extracted using the RNA Extraction Kit (AG21023, AG), and reverse transcription was performed with the Evo M-MLV RT Mix Kit (AG11728, AG). Genomic DNA was isolated using the TIANamp Genomic DNA Kit (DP304, Tiangen), and plasmid DNA was purified using the TIANprep Mini Plasmid Kit (DP103, Tiangen). Cloning procedures were carried out with the ClonExpress II One Step Cloning Kit (C112-01, Vazyme) and T4 DNA Ligase (C301-01, Vazyme). DNA fragments were gel-purified using the MolPure Gel Extraction Kit (19101ES70, Yeasen). Quantitative real-time PCR (qPCR) was conducted using ChamQ SYBR Color qPCR Master Mix (Q431-02, Vazyme). For protein analysis, total protein lysates were prepared using either Total Protein Extraction Kit (AR0103, Boster Bio) or Western and IP Cell Lysis Buffer (P0013, Beyotime), supplemented with protease and phosphatase inhibitor cocktail (P002, NCM Biotech). Protein concentrations were determined using the Bradford 1× dye reagent (5000205, Bio-Rad). Immunoprecipitation was performed using Anti-IgG Magnetic Agarose (P2171, Beyotime), Anti-DYKDDDDK Magnetic Agarose (A36798, Thermo Scientific), or Magnetic Beads-conjugated Mouse anti-DDDDK-Tag mAb (AE130, ABclonal). Recombinant His-tagged and GST-tagged proteins were purified using His-Tag Protein Purification Kit (Agarose Resin, 20751ES10, Yeasen) and GST-Tag Protein Purification Kit (20753ES10, Yeasen), respectively. Profinity IMAC Resin (uncharged, 1560121, Bio-Rad) was used for immobilized metal affinity chromatography. Mitochondrial and nuclear fractions were isolated using the Mitochondria Isolation Kit for Cultured Cells (PK10016, Proteintech) and the Nuclear Extraction Kit (PK10014, Proteintech), respectively. Cell viability was assessed using the Cell Counting Kit-8 (CCK-8, K1018, APExBIO). Mitochondrial morphology was visualized by staining with MitoTracker™ Red (M36007, Thermo Scientific). Hoechst 33342 (C1022, Beyotime) and propidium iodide (PI, 40710ES03, Yeasen) were used for nuclear and dead cell staining, respectively. Furimazine (E5806, Selleck) was used as a luciferase substrate. BeyoBlue™ Plus Coomassie Blue Ultra-Fast Staining Solution (P0004M, Beyotime) was applied for protein gel visualization. Membrane transfer for immunoblotting was performed using the eBlot L1 Rapid Transfer Kit (L00726C-75, Genscript). DTSP (57757-57-0, Solarbio) was used for chemical crosslinking, and DCEM1 (HY-171616, MCE) was used to inhibit the UPRmt. Puromycin (ant-pr-1, InvivoGen) and blasticidin S (ST018, Beyotime) were used for stable cell line selection.

Cell viability assay

Cells were seeded onto 96-well plates at a density of 1 × 10^4^ cells in 100 μL medium per well. After overnight incubation, 100 μL drug dilutions were added to the cell culture. Following the designated treatment period, 10 μL of CCK-8 reagent (K1018, APExBio) was added into each well containing 200 μL of medium. After incubation for 1 hr, the absorbance was measured at 450 nm using a Multiskan SkyHigh Microplate Spectrophotometer (Thermo Fisher Scientific). Cell viability was calculated by comparing the absorbance of cells treated with ES-Cu to that of untreated cells. In the gene functional screen, genes whose overexpression or knockdown altered cell viability by more than 30% were selected as the potential regulator of cuproptosis for further validation.

Cell death assay

Cell death was quantitatively assessed by automated high-content imaging and analysis using dual fluorescence staining with Hoechst 33342 and propidium iodide (PI). HT-1080 cells were seeded in 96-well black-walled, clear-bottom imaging plates at a density of 5×10^3^ cells per well and allowed to adhere overnight. Cells were then treated with the copper complex ES-Cu (15 nM ES plus 1.5 µM CuSO₄) or other indicated conditions. At the endpoint of treatment, cells were stained by adding Hoechst 33342 and PI directly to the culture medium at final concentrations of 5 µg/mL and 2 µg/mL, respectively, and incubated at 37°C in the dark for 30 minutes. The plate was then immediately transferred to an EVOS M7000 3D Digital Confocal Cell Imaging System (Thermo Fisher Scientific) integrated with an S Onstage 2.0 live-cell incubator, maintaining the imaging environment at 37°C with 5% CO₂. For each well, multiple non-overlapping fields of view were automatically acquired using the system’s high-resolution monochrome CMOS camera and a 10× or 20× objective, with software-driven auto-focus. Hoechst 33342 and PI fluorescence were captured sequentially using the appropriate LED light cubes and filter sets. Acquired images were processed and analyzed using the instrument’s integrated software. Nuclei were segmented based on the Hoechst 33342 signal, and each nucleus was evaluated for co-localized PI fluorescence intensity above a defined threshold. The cell death rate for each well was calculated as the percentage of PI-positive nuclei relative to the total number of Hoechst 33342-positive nuclei. Data were pooled from at least three independent experiments, each performed in technical triplicate.

Live cell imaging

For live-cell imaging, HT-1080 cells stably expressing EGFP-tagged (fused at either the N- or C-terminus) candidate genes via lentiviral transduction were used. Stable polyclonal populations were selected using blasticidin (BSD) and maintained for 1-2 passages to ensure stable expression. Cells were then seeded onto 35-mm glass-bottom confocal dishes (J40204, Jingan) in Dulbecco's Modified Eagle Medium (DMEM) supplemented with 10% fetal bovine serum (FBS) and allowed to adhere overnight.

To assess protein translocation, co-staining with organelle-specific markers was performed when necessary. For instance, to monitor mitochondrial localization, cells were incubated with 100 nM MitoTracker Red CMXRos (Beyotime, C1035) in serum-free medium for 30 minutes at 37°C prior to Hoechst 33342 staining, followed by two gentle washes with warm PBS.

Prior to imaging, cells were incubated with Hoechst 33342 (1 µg/mL) for 20 minutes at 37°C to stain nuclei, followed by a complete medium change. Time-lapse imaging was initiated immediately after the addition of the copper complex ES-Cu (15 nM ES combined with 1.5 µM CuSO₄). All imaging was performed using a Zeiss LSM 900 laser scanning confocal microscope equipped with an Airyscan 2 detector, a 40× oil-immersion objective (NA 1.4), and a stage-top incubation chamber maintaining a humidified environment at 37°C with 5% CO₂. Images were acquired sequentially using appropriate laser lines (e.g., 405 nm for Hoechst 33342, 488 nm for EGFP, 561 nm for MitoTracker Red) at defined time intervals. Image acquisition was controlled using ZEN software (blue edition, Zeiss).

Protein translocation was evaluated by both qualitative visual inspection of the time-lapse sequences and quantitative analysis of fluorescence intensity co-localization over time. Quantitative analyses, including fluorescence intensity quantification and time-course co-localization measurements, were performed using ImageJ (Fiji) with appropriate plugins (JaCoP) and GraphPad Prism 9 software.

Immunofluorescence

Cells were plated onto 35 mm confocal dishes. After incubation overnight, cells were fixed with 4% paraformaldehyde and gently agitated on a low-speed shaker for 15-20 min. Fixed cells were permeabilized with 5% (v/v) Triton X-100 for 10 min, washed three times with PBS for 5 min each and blocked with 2% BSA at RT for 30 min. Cells were incubated with anti-MRPL20 antibody (1:300, DF3663 Affinity, rabbit) diluted in Immunol Staining Dilution Buffer (P0103, Beyotime) at 4℃ overnight. After incubation, cells were washed with PBS for 5 min each. Cells were then stained with IgG-Alexa 488 (1:500, AS035, ABclonal, Donkey) diluted in PBS with 10 μg/mL Hoechst 33342 at room temperature for 1 hr. Cells were then washed three times with PBS for 5 min each. Images were acquired, and process with an LSM 900 and further analyzed by ImageJ.

Western Blot

Cells were lysed on ice for 30 minutes in cold RIPA lysis buffer (50 mM Tris-HCl, pH 7.4, 150 mM NaCl, 1% Triton X-100, 1% sodium deoxycholate, 0.1% SDS) supplemented with a protease and phosphatase inhibitor cocktail (78430, Thermo Fisher Scientific). Lysates were clarified by centrifugation at 12,000 × g for 15 minutes at 4°C. For subcellular fractionation, mitochondrial and nuclear proteins were extracted using commercial kits (Mitochondrial Extraction Kit, PK10016; Cell Nuclear Protein Extraction Kit, PK10014; both from Proteintech) strictly following the manufacturer’s protocols. The protein concentration of all lysates and fractions was determined using the Bradford assay (Bio-Rad) with bovine serum albumin as the standard.

For standard denaturing Western blot, equal amounts of protein (30-50 μg per lane) were mixed with 4× Laemmli SDS-PAGE loading buffer, boiled at 95°C for 5-10 minutes, and then separated by SDS-polyacrylamide gel electrophoresis (SDS-PAGE) on gels of appropriate acrylamide concentration. The separated proteins were electrophoretically transferred onto polyvinylidene difluoride (PVDF) membranes (Millipore) in a Tris-glycine buffer containing 20% methanol.

To analyze the native oligomeric states of proteins, a non-denaturing Native PAGE protocol was employed. Specifically, for the detection of DLAT oligomerization, proteins were extracted under native conditions using a cold, detergent-free lysis buffer (50 mM Tris-HCl, pH 7.4, 150 mM NaCl) supplemented with protease inhibitors. The lysate was centrifuged at 10,000 × g for 20 minutes at 4°C. Equal amounts of protein (20-40 μg) were mixed with a native sample buffer lacking SDS and reducing agents, loaded onto a non-denaturing polyacrylamide gel (typically 6-10% acrylamide), and electrophoresed in native running buffer (e.g., 25 mM Tris, 192 mM glycine, pH 8.3) at 4°C. Following electrophoresis, proteins were transferred to a PVDF membrane using a standard Tris-glycine transfer buffer without SDS.

For immunoblotting common to both methods, membranes were blocked for 1-2 hours at room temperature with 5% (w/v) non-fat dry milk or bovine serum albumin (BSA) in Tris-buffered saline containing 0.1% Tween-20 (TBST). Subsequently, membranes were incubated overnight at 4°C with the indicated primary antibodies. After three 10-minute washes with TBST, membranes were incubated with the corresponding horseradish peroxidase (HRP)-conjugated secondary antibodies for 1 hour at room temperature. Following another three TBST washes, protein bands were visualized using the SuperSignal West Pico PLUS chemiluminescent substrate (34577, Thermo Fisher Scientific) and imaged with a Bio-Rad ChemiDoc XRS+ imaging system. Band intensity was quantified using ImageJ (Fiji) software.

Mitochondrial Isolation

Mitochondria were isolated from cells or fresh tissues using a Mitochondria Isolation Kit (Proteintech, Cat No. PK10016) following the manufacturer’s protocol with minor optimizations. All reagents were thawed and kept on ice before use, and all centrifugation steps were performed at 4°C.

For cultured cells, approximately 2 × 10⁷ cells were harvested by centrifugation at 500 × g for 5 min, washed twice with ice-cold PBS, and resuspended in 1 mL of ice-cold Mitochondrial Isolation Reagent A supplemented with protease inhibitors. For fresh tissues, approximately 100 mg of tissue was minced on ice, washed with PBS, and homogenized in 1 mL of Reagent A. The cell or tissue suspension was transferred to a pre-chilled glass Dounce homogenizer and homogenized with 3–5 strokes (for cells) or 5–10 strokes (for tissues) on ice. Homogenization efficiency was monitored by mixing a 20 μL aliquot of the homogenate with an equal volume of 0.4% trypan blue and examining under a microscope. Homogenization was stopped when approximately 60% of cells were stained blue, indicating plasma membrane rupture.

The homogenate was carefully layered over an equal volume of Mitochondrial Isolation Reagent B in a centrifuge tube. For cell samples, the tube was centrifuged at 600 × g for 10 min; for tissue samples, at 700 × g for 10 min. The uppermost supernatant layer was collected and transferred to a new tube. This supernatant was centrifuged at 10,000 × g for 10 min to pellet the crude mitochondrial fraction.

To further purify mitochondria, the crude mitochondrial pellet was gently resuspended in 0.5 mL of Mitochondrial Wash Buffer. This suspension was then layered onto 0.5 mL of a pre-prepared high-purity mitochondrial separation medium (a 17:3 mixture of Mitochondrial Isolation Reagent C and Reagent D) in a new tube. After centrifugation at 22,000 × g for 10 min, the pellet was collected, resuspended in 1 mL of Mitochondrial Wash Buffer, and centrifuged again at 16,000 × g for 5 min. The final pellet contained highly purified mitochondria.

For validation of mitochondrial integrity and purity, an aliquot of the final mitochondrial pellet was smeared on a glass slide, stained with Janus Green B for 20 min, and examined under a light microscope. Intact mitochondria appeared as discrete blue-green rods or dumbbell-shaped structures.

Isolation of Nuclear and Cytoplasmic Proteins from Cultured Cells

Nuclear and cytoplasmic proteins were fractionated from cultured cells using the Nuclear and Cytoplasmic Protein Extraction Kit (Proteintech, Cat No. PK10014). All procedures were performed on ice or at 4°C using pre-chilled reagents and centrifuges. Approximately 2 × 10⁷ to 5 × 10⁷ cells were harvested by centrifugation at 500 × g for 5 minutes, washed twice with ice-cold PBS, and thoroughly resuspended in 1 mL of ice-cold Cytoplasmic Protein Extraction Reagent A supplemented with a protease inhibitor cocktail. The cell suspension was transferred to a pre-chilled glass Dounce homogenizer and subjected to 1–2 gentle strokes on ice to lyse the plasma membrane while preserving nuclear integrity. Lysis efficiency was monitored by mixing a 20 μL aliquot with an equal volume of 0.4% trypan blue; lysis was stopped when approximately 50% of cells were stained blue. The homogenate was incubated on ice for 10 minutes with intermittent mixing and then centrifuged at 6,500 × g for 5 minutes. The supernatant, containing the cytoplasmic protein fraction, was carefully collected. The pellet, containing intact nuclei, was washed sequentially with 1 mL of Nuclear Wash Buffer A (centrifuged at 6,500 × g for 5 minutes) and then with 1 mL of Nuclear Wash Buffer B, followed by vigorous vortexing for 1 minute (10 seconds on, 10 seconds off) and another centrifugation at 6,500 × g for 5 minutes. The washed nuclear pellet was resuspended in 100 μL of Nuclear Protein Extraction Reagent B, vortexed vigorously for 2 minutes (10 seconds on, 10 seconds off), and incubated on ice for 10 minutes. This vortex-incubation cycle was repeated twice. The suspension was then centrifuged at 16,000 × g for 10 minutes, and the resulting supernatant, containing the nuclear protein fraction, was collected. The nuclear extract was diluted 4-fold with the provided Nuclear Protein Dilution Buffer to reduce viscosity. Both cytoplasmic and nuclear protein fractions were mixed with 4× SDS-PAGE loading buffer, denatured by boiling at 100°C for 5 minutes, and stored at -80°C or used immediately for immunoblotting.

Immunoprecipitation and Specific Applications for Interaction Studies

Cells stably or transiently expressing the indicated FLAG-tagged proteins were harvested at 80-90% confluence. Cells were washed twice with ice-cold phosphate-buffered saline (PBS) and lysed on ice for 30 minutes in cold Western and IP Lysis Buffer (Beyotime, P0013) supplemented with a protease and phosphatase inhibitor cocktail (78430, Thermo Fisher Scientific). The lysates were clarified by centrifugation at 12,000 × g for 15 minutes at 4°C, and the supernatant was transferred to a new microcentrifuge tube. An aliquot of the total lysate was saved as the “Input” control.

For each immunoprecipitation reaction, 800 µg of total protein was adjusted to a volume of 500 µL with lysis buffer. To reduce non-specific binding, the lysate was pre-cleared by incubation with 30 µL of protein A/G magnetic beads for 1 hour at 4°C on a rotator. The beads were removed by magnetic separation, and the pre-cleared supernatant was transferred to a new tube.

The pre-cleared lysate was incubated with 25 µL of anti-FLAG M2 magnetic beads (M8823, Sigma-Aldrich) overnight at 4°C with gentle rotation. After incubation, the beads were collected by magnetic separation and washed three times with 1 mL of ice-cold lysis buffer. A final wash was performed with 1 mL of Tris-buffered saline (TBS, pH 7.4).

To elute the bound proteins, the beads were resuspended in 60 µL of 2× SDS-PAGE loading buffer containing 100 mM DTT. The mixture was boiled at 95°C for 10 minutes to denature the proteins.

The eluted proteins (IP eluate), along with the Input samples, were separated by SDS-PAGE and analyzed by Western blotting using antibodies against the FLAG tag (for IP validation) and the HA tag (for detecting co-precipitated proteins), as described in the Western Blot Analysis section.

To investigate OXA1L self-dimerization and its interaction with Tim23, HT-1080 cells were co-transfected with plasmids encoding OXA1L-HA and OXA1L-FLAG, or with OXA1L-HA and Tim23-FLAG, respectively. FLAG immunoprecipitation was then performed as described above. Successful co-immunoprecipitation of the HA-tagged protein with the FLAG-tagged bait confirmed the specific protein-protein interaction.

NanoLuc Complementation Assay

The NanoLuc complementation assay was performed using constructs in which the coding sequences of OXA1L, its triple mutant OXA1L-Mut (H170A/H172A/H209A), and candidate mitochondrial ribosomal proteins (MRPL20, MRPL13, MRPL28, MRPL48, MRPL49, MRPL51) as well as TIM23, were cloned into the split NanoLuc vector backbone described in the previous section. Specifically, OXA1L and OXA1L-Mut were fused to either fragment 1 (F1) or fragment 2 (F2) of NanoLuc. All other candidate interactors were fused to the F2 fragment. An unfused F2 fragment construct served as the baseline negative control.

For the assay, HT-1080 cells were seeded in 24-well plates and co-transfected with the indicated pairs of plasmids using Lipofectamine 3000. To investigate OXA1L self-association, cells were co-transfected with OXA1L-F1 and either OXA1L-F2 or the empty F2 control. To assess the impact of the triple mutation on self-association, cells were co-transfected with OXA1L-Mut-F1 and either OXA1L-F2 or the F2 control. To screen for interactions with candidate proteins, cells were co-transfected with either OXA1L-F1 or OXA1L-Mut-F1 and each of the F2-tagged candidate proteins.

Twenty-four hours post-transfection, cells were treated with the copper complex ES-Cu (1.5 µM CuSO₄ and 15 nM Elesclomol) for 10 hours or left untreated as a control. Following treatment, luminescence was measured using the Nano-Glo Live Cell Assay System (Promega). The NanoLuc substrate was added directly to the culture medium, incubated for 5 minutes at room temperature, and the luminescence signal was recorded immediately using a microplate reader.

The strength of protein-protein interaction was quantified by calculating a Normalized Luminescence Ratio. For each tested pair (A and B), the NLR was defined as: NLR = Luminescence (A–B) / [Luminescence (A alone) + Luminescence (B alone)], where “A–B” represents the luminescence from cells co-expressing the F1-fusion protein A and the F2-fusion protein B, and “A alone” or “B alone” represent the background signals from cells expressing the F1- or F2-fusion protein paired with the corresponding empty vector control. All experiments included three biological replicates. The resulting NLR values were compared across treatment conditions and visualized as heatmaps for analysis.

Detection of OXA1L Dimerization by DTSP Cross-linking

Purified WT-OXA1L and Mut-OXA1L proteins (14 μM) were incubated at 25°C for 1 hour in a buffer containing 20 mM HEPES-KOH (pH 7.6), 2 mM MgCl₂, and 100 mM KCl under the following conditions: untreated control, 10 μM ES-Cu treatment, and co-treatment with 10 μM ES-Cu and 10 μM TTM. Subsequently, DTSP was added to a final concentration of 1 mM to initiate cross‑linking. After 30 minutes, the reaction was quenched with 50 mM Tris-HCl (pH 7.6). Samples were processed under non‑reducing conditions and analyzed by 15% SDS-PAGE followed by Western blotting using an anti-OXA1L antibody to detect dimer bands at approximately 90 kDa.

Analysis of OXA1L High–Molecular–Weight Complexes and Cleavage Fragments

To analyze the native oligomeric state of OXA1L and detect potential copper-induced cleavage fragments, we performed experiments in both wild-type and YME1L-knockout genetic backgrounds. HEK293T cells (wild-type or stably expressing YME1L-targeting sgRNA) were transiently transfected with plasmids encoding double-tagged OXA1L (FLAG-OXA1L-HA) or its copper-binding mutant (FLAG-OXA1L-Mut-HA, H170A/H172A/H209A). At 24 hours post-transfection, cells were treated with the copper complex ES-Cu (1.5 μM CuSO₄ and 15 nM Elesclomol) for 10 hours or left untreated as a control. Following treatment, cells were harvested and mitochondria were isolated using a Mitochondria Isolation Kit (Proteintech, Cat No. PK10016). The purified mitochondrial pellet was resuspended in Mitochondria Storage Buffer (Beyotime, C3609) and incubated at 37°C for 1 hour to preserve native protein complexes.

To solubilize mitochondrial complexes under non-denaturing conditions, the mitochondrial suspension was lysed in ice-cold lysis buffer containing 1% (w/v) digitonin, 20 mM HEPES (pH 7.4), 100 mM NaCl, and 2 mM MgCl₂, supplemented with a protease inhibitor cocktail, for 30 minutes on ice. The lysate was clarified by centrifugation at 20,000 × g for 20 minutes at 4°C. To specifically enrich for OXA1L-containing complexes, the clarified lysate was incubated with anti-FLAG M2 magnetic beads for 2 hours at 4°C with gentle rotation. After extensive washing with lysis buffer containing 0.1% digitonin, bound complexes were competitively eluted by incubating the beads with 3× FLAG peptide (150 ng/μL) for 30 minutes at 4°C.

The eluted proteins were mixed with native PAGE sample buffer and separated on a 4–20% gradient Bis-Tris polyacrylamide gel under native conditions. For comparison, an aliquot of the same eluate was denatured and resolved on a 4–20% gradient SDS-polyacrylamide gel. Proteins were transferred to PVDF membranes. To avoid potential detection interference from the competing FLAG peptide, immunoblotting was performed using a monoclonal anti-HA antibody. This approach enabled the specific visualization of HA-tagged OXA1L species, allowing for comparative analysis of its distribution within high-molecular-weight complexes and the accumulation of cleavage fragments, in response to ES-Cu treatment, across both wild-type and YME1L-deficient cellular contexts.

Purification of Recombinant His-Tagged Proteins

The coding sequences of human ATF5, DNAJA3, HSPA9, HSPD1, and HSPE1 were cloned into the pET-28a (+) vector for expression with an N-terminal 6×His tag. Each construct was transformed into E. coli BL21(DE3) competent cells. A single colony was inoculated into LB medium containing 50 µg/mL kanamycin and grown overnight at 37°C. The overnight culture was diluted 1:100 into fresh LB medium and grown at 37°C until the OD600 reached 0.6-0.8. Protein expression was induced by adding isopropyl β-D-1-thiogalactopyranoside (IPTG) to a final concentration of 0.5 mM, followed by incubation at 16°C for 16-18 hours. Cells were harvested by centrifugation at 6,000 × g for 15 minutes at 4°C.

The cell pellets were resuspended in lysis buffer (20 mM Tris-HCl, pH 8.0, 300 mM NaCl, 10 mM imidazole) supplemented with 1 mM phenylmethylsulfonyl fluoride (PMSF) and lysed by sonication on ice. The lysate was clarified by centrifugation at 15,000 × g for 30 minutes at 4°C. The supernatant was incubated with HisSep Ni-NTA Agarose Resin (Yeasen, 20502ES80) for 1 hour at 4°C with gentle agitation. The resin was washed with 10 column volumes of wash buffer (20 mM Tris-HCl, pH 8.0, 300 mM NaCl, 30 mM imidazole). The bound His-tagged proteins were eluted with elution buffer (20 mM Tris-HCl, pH 8.0, 300 mM NaCl, 300 mM imidazole). The eluted proteins were dialyzed against storage buffer (20 mM Tris-HCl, pH 7.4, 150 mM NaCl, 10% glycerol) to remove imidazole. Protein concentration was determined using the Bradford assay, and purity was assessed by Coomassie Brilliant Blue staining following SDS-PAGE.

In Vitro His Pull-Down Assay

To investigate direct protein-protein interactions, an in vitro pull-down assay was performed using purified His-tagged proteins as the immobilized bait. For each reaction, 10 µg of a purified His-tagged protein (His-ATF5, His-DNAJA3, His-HSPA9, His-HSPD1, or His-HSPE1) was immobilized on 20 µL of HisSep Ni-NTA Agarose Resin (Yeasen, 20502ES80) by incubating in binding buffer (20 mM Tris-HCl, pH 7.4, 150 mM NaCl, 0.1% Triton X-100, 10 mM imidazole) for 2 hours at 4°C. Resin incubated with binding buffer alone served as a negative control.

After immobilization, the resin was washed three times with the same binding buffer to remove unbound protein. Then, 10 µg of the commercially purchased recombinant human GST-MRPL20 protein was added to the His-tagged bait-bound resin in 500 µL of binding buffer and incubated overnight at 4°C with gentle rotation. Following incubation, the resin was collected by brief centrifugation and washed five times with 1 mL of ice-cold binding buffer. Bound proteins were competitively eluted by incubating the resin with 40 µL of elution buffer (20 mM Tris-HCl, pH 7.4, 150 mM NaCl, 300 mM imidazole) for 30 minutes at 4°C. The eluate was mixed with 5× SDS-PAGE loading buffer and denatured. The input proteins (5% of bait and prey), the unbound flow-through, and the eluted samples were separated by SDS-PAGE and transferred to PVDF membranes. The interaction was assessed by Western blotting using an anti-GST antibody to detect the pulled-down GST-MRPL20 protein.

Metal Binding Assay

This assay was performed using Profinity IMAC resins (Bio-Rad) charged with specific metal ions. The bead preparation protocol involved an initial wash cycle (five repeats) with ultra-pure H₂O. The clean resins were then coated by incubation in 0.2 M solutions of the designated metal salts for 2 hours at 4°C. Excess metal was removed through subsequent washing steps: five washes with H₂O and two washes with acidic buffer (0.05 M sodium acetate, 0.5 M NaCl, pH 4.0). The metal-coated beads were finally stored at 4°C in binding buffer (0.05 M sodium phosphate, 0.5 M NaCl, pH 8.0). For the binding experiment, cells were lysed in ice-cold NTN buffer (100 mM NaCl, 20 mM Tris–HCl pH 8.0, 0.5% Nonidet P-40) containing a protease/phosphatase inhibitor cocktail. The resulting lysates were mixed with the prepared metal beads and incubated overnight at 4°C with gentle agitation.

RNA Isolation and Quantitative Real-Time PCR (qRT-PCR)

Total RNA was isolated from HT-1080 cells using RNAex Pro RNA isolation reagent (AG21101 & AG21024, AG) according to the manufacturer’s instructions. Briefly, cells were lysed directly in culture dishes with the reagent, and the lysate was transferred to a phase-separation tube. Following the addition of chloroform and vigorous mixing, the aqueous phase containing RNA was separated by centrifugation. The RNA was then precipitated with isopropanol, washed with 75% ethanol, and finally dissolved in RNase-free water. RNA concentration and purity were assessed by measuring the absorbance at 260 nm and 280 nm using a spectrophotometer (Nanodrop). The A260/A280 ratios of all samples were between 1.8 and 2.0, indicating high purity. RNA integrity was confirmed by visualizing the distinct 18S and 28S ribosomal RNA bands on a 1% agarose gel.

For cDNA synthesis, 1 µg of total RNA was used as the template. Genomic DNA (gDNA) removal and reverse transcription were performed sequentially in a single tube using the EVO M-MLV Reverse Transcription Kit (AG11728, AG). The reaction was assembled according to the kit protocol. Briefly, RNA was first incubated with the provided gDNA Remover at 42°C for 2 minutes to eliminate genomic DNA contamination. Subsequently, reverse transcription was carried out by adding the reverse transcriptase and reaction mix, followed by incubation under the following conditions: 37°C for 15 minutes (reverse transcription), 85°C for 5 seconds (enzyme inactivation), and a final hold at 4°C. The synthesized cDNA was diluted 5-10 fold with nuclease-free water and stored at -20°C until use.

Quantitative real-time PCR (qRT-PCR) was performed on a CFX Connect Real-Time PCR Detection System (Bio-Rad) using the 2× SYBR Green Pro Taq HS Premix (AG11733, AG). Each reaction was conducted in a 20 µL final volume containing 10 µL of 2× SYBR Green mix, 0.8 µL (0.4 µM final concentration) of each gene-specific forward and reverse primer (primer sequences are listed in Table S8), 2 µL of diluted cDNA template, and 7.2 µL of nuclease-free water. The thermal cycling protocol consisted of an initial denaturation at 95°C for 30 seconds, followed by 40 cycles of denaturation at 95°C for 5 seconds and annealing/extension at 60°C for 30 seconds. A melt curve analysis was performed at the end of each run from 65°C to 95°C with an increment of 0.5°C every 5 seconds to confirm the specificity of the amplification. All reactions were performed in technical triplicates, and no-template controls (NTC) were included in each run to rule out contamination. The relative expression level of each target gene was calculated using the 2−ΔΔCt method, with GAPDH serving as the endogenous reference gene for normalization.

Spatio-Temporal Mass Spectrometry in living cells

HT1080 cells transiently transfected with miniTurboID plasmids were treated with 1.5 μM CuSO_4_ and 15 nM ES and harvested at a series of time points 0, 2, 4, 6, 8, and 10 hrs. 500 μM biotin (HY-B0511, MCE) was added into the medium 20 min prior to collection. After thoroughly washing with ice-cold PBS three times, cells were lysed in the lysis buffer (25 mM Tris, pH 7.2, 150 mM NaCl, 0.8% Triton X-100, and 1x protease inhibitor). Cell lysates were sonicated at 4℃, with three bursts of 10 s with an interval of 2 s between bursts. Cell debris was removed by centrifuged at 16,000 g for 20 min. The supernatant was then mixed with pre-washed Pierce™ streptavidin Magnetic Agarose Beads (78609, Thermo Fisher Scientific) and gently agitated on a rocking platform at 4℃ for 2.5 hrs.

After incubation, the Streptavidin Agarose Resin was centrifuged at 500 g for 1 min, and the supernatant was removed. The resin was subjected to two washes with lysis buffer and four washes with PBS with each wash followed by centrifugation at 500 g for 1 min. 40 μL loading buffer (100 mM Tris-HCl, 4% SDS, 1 mM DTT, pH 7.6) was added to Streptavidin Agarose Resin, and the resin was then boiled at 100℃ for 20 min. After centrifugation at 14,000 g for 20 min to separate precipitation and supernatant, the supernatant was collected and stored at -80℃.

Protein quality was detected by Bradford protein quantitative method and 12% SDS-PAGE gel electrophoresis with coomassie brilliant blue staining. For quantitative proteomics studies, protein lysates were extracted using lysis buffer (8 M Urea, 100 mM TEAB, pH 8.5). The lysates were then reduced with 10 mM TCEP and alkylated with 50 mM iodoactamide, followed by acetone precipitation. The precipitated proteins were resuspended in 100mM TEAB and digested overnight at 37℃ with Trypsin (V5280, Promega). The resulting peptides are desalted using C18 desalting column (89852, Thermo Fisher Scientific). After three washes with a buffer containing 0.1% formic acid and 3% acetonitrile, the peptides were eluted with a buffer of 0.1% formic acid and 70% acetonitrile, and then lyophilized.

Immunoprecipitation-mass Spectrometry Data Pre-processing and Normalization

Raw protein intensities were first filtered to remove common contaminants (e.g., keratins, immunoglobulins) and low-confidence identifications. Proteins were retained only if they contained at least 2 unique peptides and were detected in at least 2 biological replicates within at least one experimental group (Control or ES-Cu). Filtered intensities were log2-transformed. To correct for variations in immunoprecipitation efficiency, a bait-normalization strategy was applied: the log2 intensity of the bait protein (MRPL20) was subtracted from all other proteins in the corresponding sample, followed by re-centering to the median bait intensity across all samples. Missing values were assumed to be missing not at random (MNAR) and were imputed using a left-censored normal distribution (MinProb). Data quality was assessed at multiple stages: sample relationships were evaluated using Principal Component Analysis (PCA) and hierarchical clustering, and immunoprecipitation efficiency was validated by monitoring the abundance rank of the bait protein (MRPL20) across all samples.

Differential protein-protein interactions were identified using the Limma package in R. To account for potential heteroscedasticity and varying sample quality, a linear model was fitted for each protein with sample-specific quality weights estimated via the arrayWeights function. Comparisons of interest (ES-Cu vs. Control) were defined using a contrast matrix, and empirical Bayes statistics were applied to moderate t-statistics. P-values were adjusted for multiple testing using the Benjamini-Hochberg (BH) method. Significant interactions were classified as "Strong" hits (Log2FC > 1, FDR < 0.05) or "Moderate" hits (Log2FC > 0.58, FDR < 0.10). A sensitivity analysis was performed to ensure that the identification of key interactors was not biased by individual sample variations. Gene Ontology (GO) enrichment analysis was performed using the clusterProfiler package. The list of strongly upregulated, differential proteins (Log2FC > 1, FDR < 0.05) was used as the gene set of interest, while the complete list of filtered, imputed proteins (n = 2,780) served as the statistical background (universe) to avoid selection bias. Over-Representation Analysis (ORA) was conducted using the enrichGO function for Biological Process (BP), and Cellular Component (CC) ontologies. P-values were adjusted using the BH method, and terms with an adjusted P-value < 0.05 were considered significantly enriched.

Bioinformatics Analysis

Gene expression analysis of UPR^mt^ components and copper death-related genes was performed using the GEPIA 3 web server (https://gepia3.bioinfoliu.com/), which integrates RNA-Seq data from the TCGA and GTEx projects using a standard processing pipeline. For pan-cancer analysis, differential expression was assessed across 15 major cancer types, and the Log2 Fold Change (Tumor/Normal) was visualized as a heatmap using the pheatmap package in R. For the COAD-specific analysis, the median expression levels (Log2 TPM + 1) of genes of interest in tumor and normal tissues were extracted, and statistical significance was determined by the web server using t-test. The results were visualized as grouped bar plots using the ggplot2 package.

Bioinformatic Prediction of Cu²⁺-Binding Sites

To identify putative copper-binding motifs in OXA1L, we conducted an in silico analysis using the MIB2 (Metal Ion-Binding site prediction server, version 2). This server predicts metal-binding residues through a local structure comparison algorithm. It utilizes a library of experimentally verified metal-binding structural templates (within a 3.5 Å sphere of the metal ion) and aligns them to the query protein structure using a fragment transformation approach. A composite scoring function, integrating sequence and structural conservation, is then used to assess the alignment. The high-confidence AlphaFold2-predicted structure of human OXA1L (Q15070) was submitted to the server. The analysis was configured to predict binding sites specifically for divalent copper ions (Cu²⁺). All predicted residues with scores exceeding the default statistical threshold, as determined by MIB2’s internal benchmarking, were reported and considered for subsequent experimental validation.

Molecular Docking Simulation

To model the interaction geometry and binding affinity between Cu²⁺ and the predicted residue cluster, molecular docking was performed directly within the MIB2 server framework. The server utilizes the predicted binding site as the target pocket for docking calculations. The Cu²⁺ ion was treated as a rigid body, and its optimal positioning within the constraints of the pre-defined binding residue cluster was computationally sampled. The docking algorithm evaluates multiple poses and outputs the most energetically favorable conformation based on scoring functions that account for electrostatic interactions, van der Waals forces, and geometric complementarity. The resulting model, depicting the coordination of Cu²⁺ by the specific side chains of OXA1L, was visualized and analyzed.

Mass spectrometry workflow and data analysis

Mass spectrometry (MS)-based proteomics were analyzed by using the Parallel Accumulation Serial Fragmentation (PASEF®) technology with timsTOF Pro (Burker, America) at Novogene (China). 2D-LC-MS is performed on peptides, where in the first dimension of chromatographic analysis, SCX or high pH HPLC is primarily used to preliminarily separate the peptide mixture into multiple fractions. This step reduces the complexity of the peptide sample. Each of these fractions is then subjected to LC-MS to achieve better mass spectrometry identification results. Whole scanning range of MS was divided into several Windows, and all ions in each window were selected, broken and detected in a cycle.

The quantification type chosen is DIA (Data Independent Acquisition), Spectronaut-Pulsar is used to search the database and build the database. The parameters are: Mass tolerance for precursor ion was set to 10 parts per million (ppm), and for the product ion, it was set to 0.02 Daltons (Da). Carbamidomethylation of cystein was designated as fixed modification, Oxidation of methionine (M) was considered a variable modification. Acetylation was specified as a modification occurring at the N-terminus. Further screening using Spectronaut-Pulsar retained peptides with more than 99% confidence. Import the DIA data into the Spectronaut (Biognosys) software and extract ion pairs of chromatographic peaks. Calculate the peak area and achieve the qualitative and quantitative of peptides.

Proteomics expression data preprocessing

Quantitative proteomics data preprocessing was conducted in R (v4.4.2) using a stratified quality control framework. For each organelle-time point combination (6 time points × 5 organelles), proteins were sequentially filtered based on the following criteria: (1) more than one missing value in the triplicate measurements; (2) detection in fewer than two non-zero biological replicates; (3) technical variability exceeding the 90th percentile of condition-specific coefficients of variation; and (4) mean abundance below the 10th percentile within their organelle-time group. After filtering, remaining missing values were imputed using condition-specific row means (excluding zeros when applicable). To achieve gene-level resolution, the most abundant protein isoform for each gene was retained. Filtered protein intensity values were then log2-transformed using a zero-preserving approach, where only non-zero values were transformed, while zero values were maintained as zeros. This transformation was performed independently for each organelle to retain the zero-coded entries for undetected measurements; these entries do not establish biological absence. Following transformation, datasets from individual organelles were merged into a comprehensive matrix. Zeros were preserved for proteins not detected in certain organelles. Finally, the mean intensity of each protein was calculated for each organelle–time point combination, resulting in a dataset of 2,867 high-confidence proteins with organelle-specific temporal profiles.

Spatial-temporal proteomics expression pattern definition

To define spatiotemporal expression patterns, the preprocessed mean intensity of each protein was standardized using Z-score normalization. Clustering and heatmap visualization of the proteomics expression data were performed using the ComplexHeatmap package (v2.18.0), applying the Ward.D2 clustering method and Euclidean distance as the distance metric. Proteins were divided into ten clusters based on their expression patterns across organelles and time points by setting the row_km parameter to 10. For functional characterization, clusterProfiler (v4.10.0) was used to perform GO enrichment analysis on the genes within each cluster, with the background gene set restricted to GO Molecular Function. Pathways with p.adjust < 0.1 and FoldEnrichment > 1.5 were retained and sorted by FoldEnrichment. The resulting pathways were visualized to highlight the functional characteristics of each cluster, with the top 30 pathways shown for clusters enriched with more pathways.

Differential protein expression analysis at each organelle

Differential expression analysis was conducted using the limma package (v3.60.6) on organelle-specific, zero-preserving log2-transformed data. For each organelle, protein abundance at experimental time points (2, 4, 6, 8, and 10 hours) was compared to the baseline (0 hours) using linear models with empirical Bayes moderation (trend=TRUE to account for intensity-dependent variance). Proteins with no detectable expression across all samples within a given organelle–time point comparison were excluded. Adjusted p-values were calculated using the Benjamini-Hochberg method to control the false discovery rate (FDR). Proteins were classified as differentially expressed if they met the thresholds of adjusted p-value < 0.01 and |log2 fold change| > 2. This analysis identified high-confidence differentially expressed proteins (DEPs) with distinct organelle-specific temporal expression patterns. Results were visualized using volcano plots, with color-coded markers highlighting significantly upregulated and downregulated proteins.

Subcellular proteome trafficking analysis

In this analysis, organelle localization at each time point was determined using a maximum abundance-based method applied to the preprocessed quantitative proteomics data. Protein abundance data for each organelle and time point were extracted, and the coefficient of variation (CV) of replicate measurements was calculated. Proteins with a CV ≤ 0.3 were retained to ensure data reliability. Abundance values were then median-normalized across replicates to eliminate experimental biases. For each protein, normalized abundance values across organelles were ranked to identify the maximum abundance (Max_Abundance) and the second-highest abundance (Second_Max_Abundance). A confidence score was calculated as Confidence_Score = Max_Abundance / (Second_Max_Abundance + 0.01) to quantify the specificity of localization. Proteins were classified into three categories based on predefined thresholds: Unique (Confidence_Score > 10), Multiple (2 < Confidence_Score ≤ 10), and Uncertain (Confidence_Score ≤ 2). The organelle with the highest abundance was assigned as the final localization for each protein.

Proteins classified as Unique were further compared with known organelle annotations from publicly available databases, including Uniprot, The Human Protein Atlas, and Gene Ontology Cellular Component (GO CC), to evaluate the consistency of predicted localizations with existing knowledge. The distribution of protein assignment types (Unique, Multiple, and Uncertain) was summarized and visualized across all time points to assess temporal dynamics in organelle localization. To illustrate subcellular trafficking patterns, the movement of proteins among five organelles across six time points was visualized using a Sankey diagram generated with the ggalluvial package (v0.12.5) and ggplot2 package (v3.5.1).

Xenograft Model and In Vivo Treatment

All animal procedures were conducted in strict accordance with relevant ethical regulations and were approved by the Institutional Animal Care and Use Committee of Zhejiang University (Protocol No. ZJU20250707). Four-week-old female, specific pathogen-free (SPF) BALB/c nude mice were obtained from the Laboratory Animal Center of Zhejiang University and housed under standard conditions with ad libitum access to food and water. For tumor establishment, HCT-116 cells (6 × 10⁶ cells suspended in a 1:1 mixture of Matrigel and PBS) were subcutaneously injected into the right flank of five-week-old female mice. Tumor growth was monitored using digital calipers, and tumor volume was calculated with the formula: Volume = (Length × Width²) / 2.

The in vivo study was conducted in two sequential phases. In the first phase, aimed at determining a safe and sub-therapeutic dose of DCEM1, mice bearing tumors (50-100 mm³) were randomly assigned to receive vehicle control (DMSO) or DCEM1 at 10, 20, or 40 mg/kg via intraperitoneal (i.p.) injection every other day (n = 6 per group). Based on the results indicating that 20 mg/kg DCEM1 was well-tolerated and did not significantly inhibit tumor growth as a single agent, this dose was selected for the subsequent combination therapy phase.

In the second phase, a new cohort of tumor-bearing mice was randomized into the following five treatment groups (n = 6 per group) when tumor volumes reached 50-100 mm³: (1) Vehicle control (DMSO); (2) ES-Cu (ES: 20 mg/kg + CuSO₄: 0.5 mg/kg); (3) DCEM1 (20 mg/kg); (4) ES-Cu + DCEM1 (20 mg/kg); (5) ES-Cu + DCEM1 (20 mg/kg) + TTM (10 mg/kg). All agents were administered via i.p. injection every other day.

Throughout both experimental phases, mouse body weight and tumor volume were recorded every other day, coinciding with treatment administration. The maximum tumor burden permitted was 1500 mm³, which was not exceeded. At the experimental endpoint, mice were euthanized, and tumors were excised, weighed, and processed for further analysis.

Statistical Analysis

Statistical analysis was performed using GraphPad Prism (8.3.1). All experimental data shown in plots are represented as mean ± SD/SEM, and exact number of independent replicates for each experiment is stated in figure legends. Significant differences were determined by one-way ANOVA or two-tailed t-tests. The significance level between two groups was set to 0.05 and the exact p-values were shown in the figures.

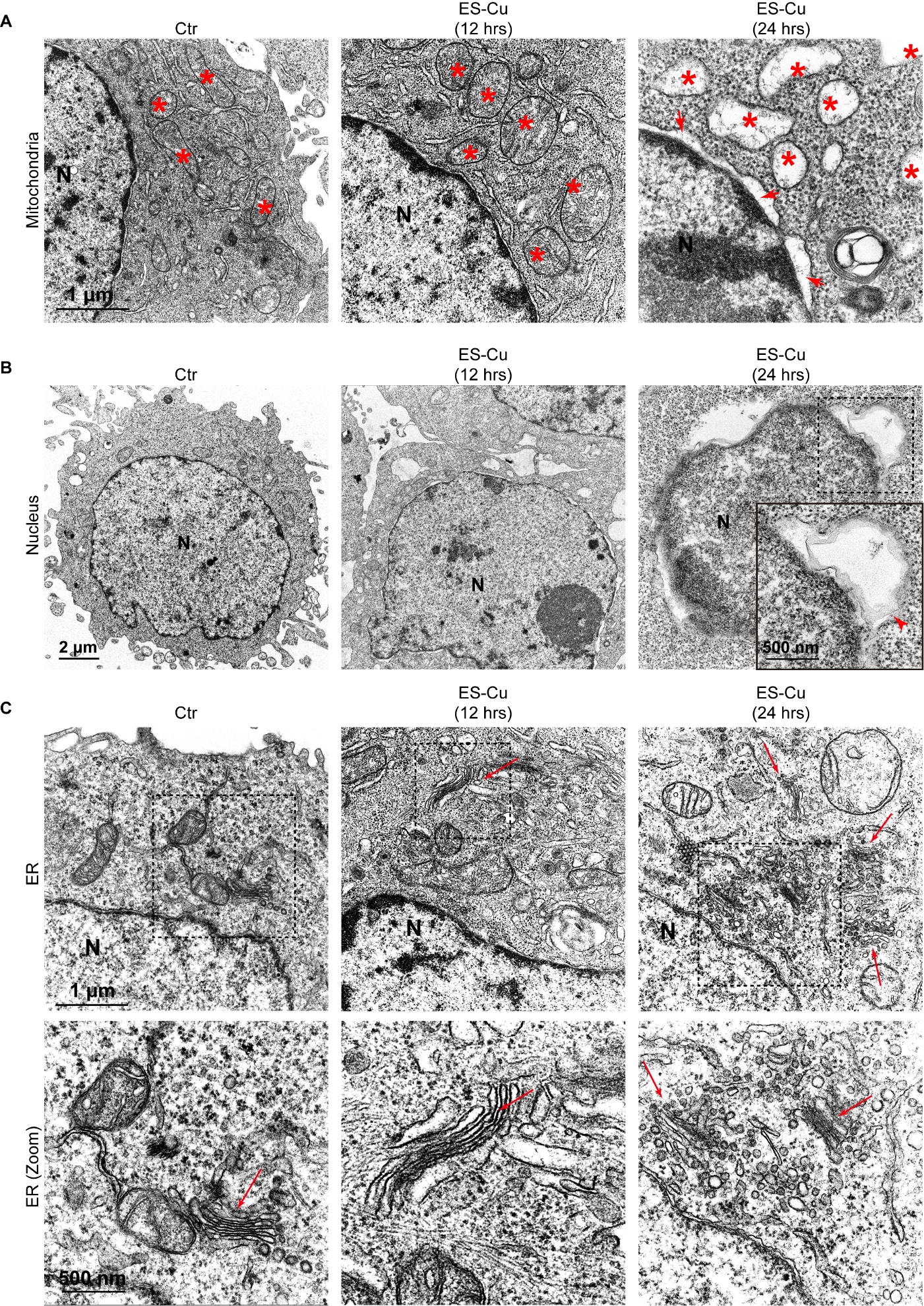

Fig. S1. ES–Cu induces progressive ultrastructural changes in mitochondria, nuclei and the endoplasmic reticulum.

(A–C) Representative transmission electron micrographs under control conditions or following ES–Cu treatment for 12 or 24 h. (A) Mitochondrial morphology at the indicated times; red asterisks mark mitochondria. (B) Nuclear morphology and changes at the nuclear boundary; an enlarged region of the 24-h sample is shown in the inset. (C) Endoplasmic reticulum (ER) morphology and higher-magnification views of the boxed regions; arrows mark the indicated ER structures. Scale bars are shown in the images.

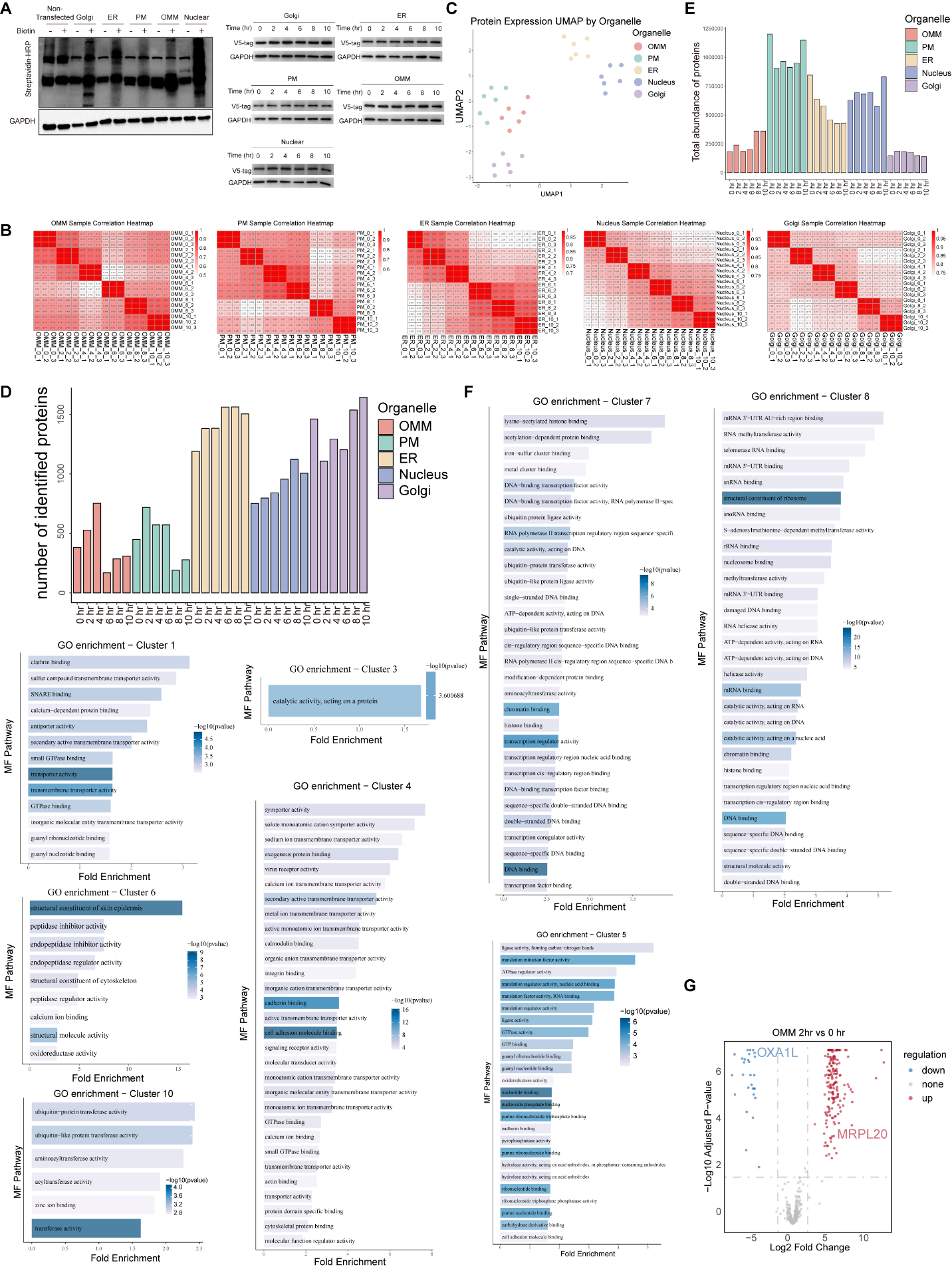

Fig. S2. Quality control and functional annotation of the spatiotemporal proteomic atlas during copper stress.

(A) Validation of organelle-targeted miniTurbo labeling in HT-1080 cells. Streptavidin–HRP detects biotinylated proteins in untransfected and organelle-targeted miniTurbo-expressing cells with or without 500 μM biotin (left); anti-V5 immunoblots show the indicated constructs over 0–10 h of ES–Cu treatment (right). GAPDH is the loading control. (B) Pairwise Pearson correlation heatmaps of three proteomic replicates per time point for outer mitochondrial membrane (OMM), plasma membrane (PM), ER, nucleus and Golgi samples. (C) Uniform manifold approximation and projection (UMAP) of the proteomic samples by organelle. (D) Numbers of quantified proteins in the biotinylated pool for each compartment at 0, 2, 4, 6, 8 and 10 h. (E) Summed protein intensity in the biotinylated pool by compartment and time point. Panels (D) and (E) describe the streptavidin-enriched fraction rather than total organelle protein content. (F) Gene Ontology molecular-function enrichment for clusters 1, 3, 4, 5, 6, 7, 8 and 10 from Fig. 1D. Terms met adjusted P < 0.1 and fold enrichment > 1.5; clusters 2 and 9 had no terms meeting these criteria. (G) Volcano plot of OMM proteins at 2 h relative to 0 h, highlighting OXA1L and MRPL20.

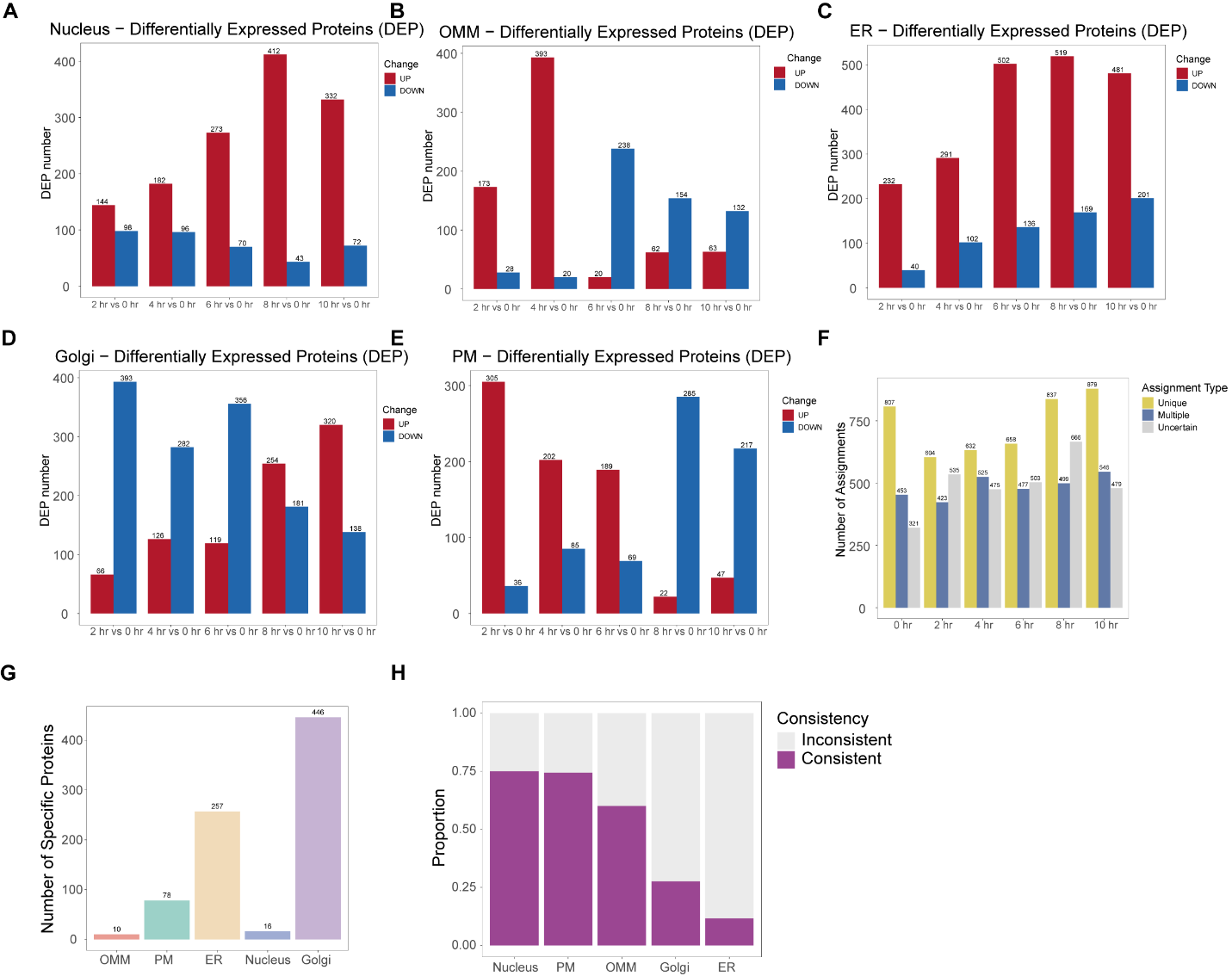

Fig. S3. Organelle-specific proteomic responses and localization assignments over the copper-stress time course.

(A–E) Numbers of upregulated (red) and downregulated (blue) proteins at 2, 4, 6, 8 and 10 h versus 0 h in the nucleus (A), OMM (B), ER (C), Golgi (D) and PM (E). (F) Distribution of protein localization assignments classified as Unique, Multiple or Uncertain across the time course. (G) Numbers of proteins assigned uniquely to each compartment at baseline (0 h). (H) Fractions of uniquely assigned proteins whose baseline localization agrees or disagrees with publicly available localization annotations. See Materials and Methods for the assignment criteria.

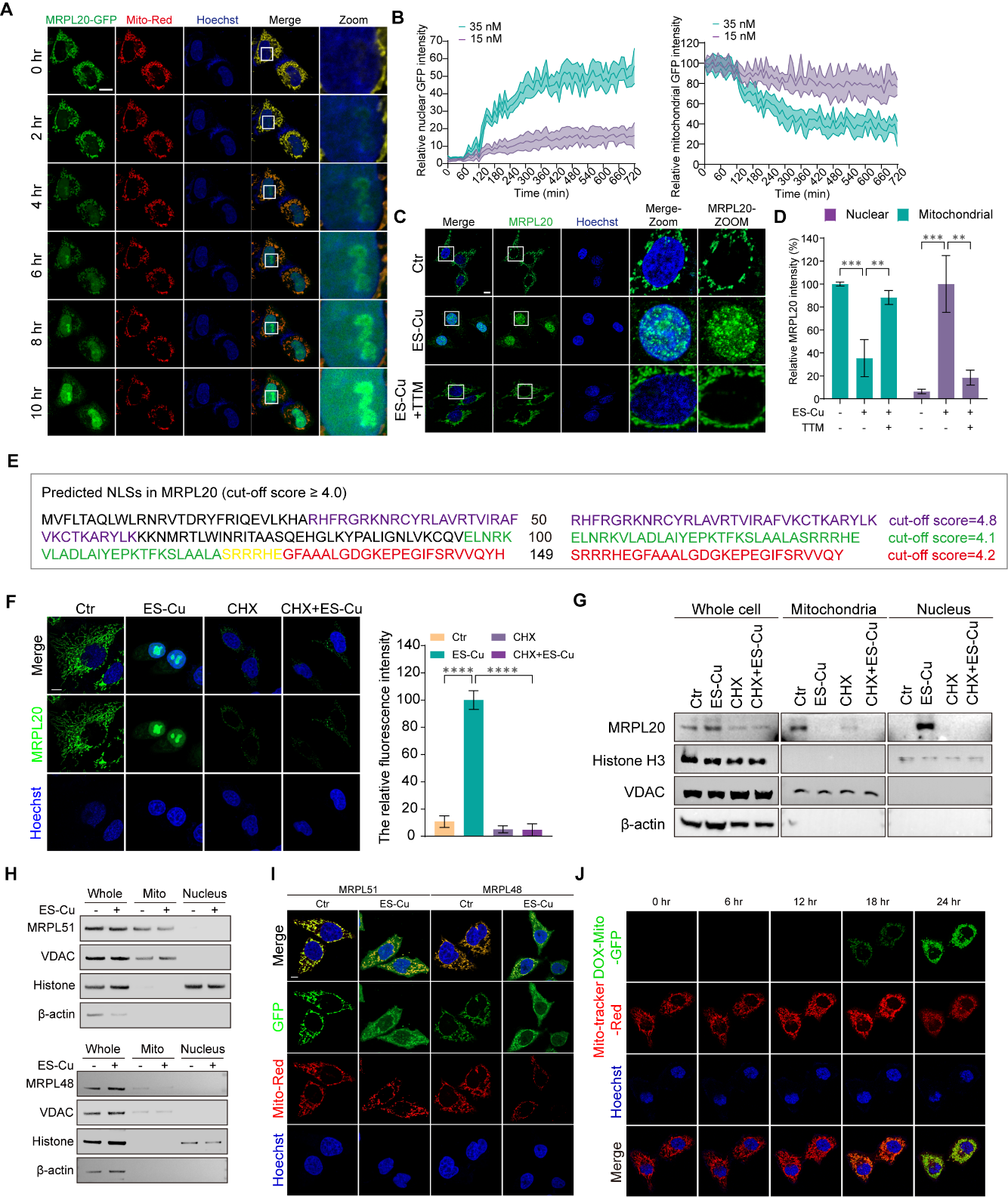

Fig. S4. Copper stress induces selective nuclear redistribution of MRPL20.

(A) Time-lapse confocal images of MRPL20–GFP (green), Mito-Red (red) and Hoechst (blue) in HT-1080 cells during ES–Cu exposure (1.5 μM CuSO₄ plus 35 nM elesclomol). Boxed nuclear regions are enlarged at right; scale bar, 10 μm. (B) Time courses of GFP fluorescence in cells treated with 1.5 μM CuSO₄ plus 15 or 35 nM elesclomol; lines show the mean and shaded variability (n = 6). (C) Immunofluorescence of endogenous MRPL20 (green) and Hoechst-stained nuclei (blue) following ES–Cu (1.5 μM CuSO₄ plus 15 nM elesclomol, 12 h), with or without 20 μM tetrathiomolybdate (TTM). Boxed regions are enlarged at right; scale bar, 10 μm. (D) Quantification of mitochondrial and nuclear MRPL20 fluorescence for the conditions in (C), normalized to untreated mitochondrial MRPL20 (mean ± SEM, n = 6). (E) Putative intrinsic MRPL20 nuclear localization sequences predicted by cNLS Mapper; the indicated sequence segments meet the displayed cutoff score. Residues shown in yellow belong to both the green and red predicted NLS motifs. (F) MRPL20 immunofluorescence after cycloheximide (CHX) pretreatment with or without ES–Cu. (G) MRPL20 immunoblots of whole-cell, mitochondrial and nuclear fractions for the conditions in (F); histone H3, VDAC and β-actin are compartment and loading controls. (H) Immunoblots of MRPL51 and MRPL48 in whole-cell, mitochondrial and nuclear fractions with or without ES–Cu. (I) Images of MRPL51- and MRPL48-associated GFP fluorescence, Mito-Red and Hoechst in control and ES–Cu-treated cells. (J) Time-course images of doxycycline-inducible mitochondrially targeted GFP (green), MitoTracker Red (red) and Hoechst (blue) at 0, 6, 12, 18 and 24 h of ES–Cu exposure.

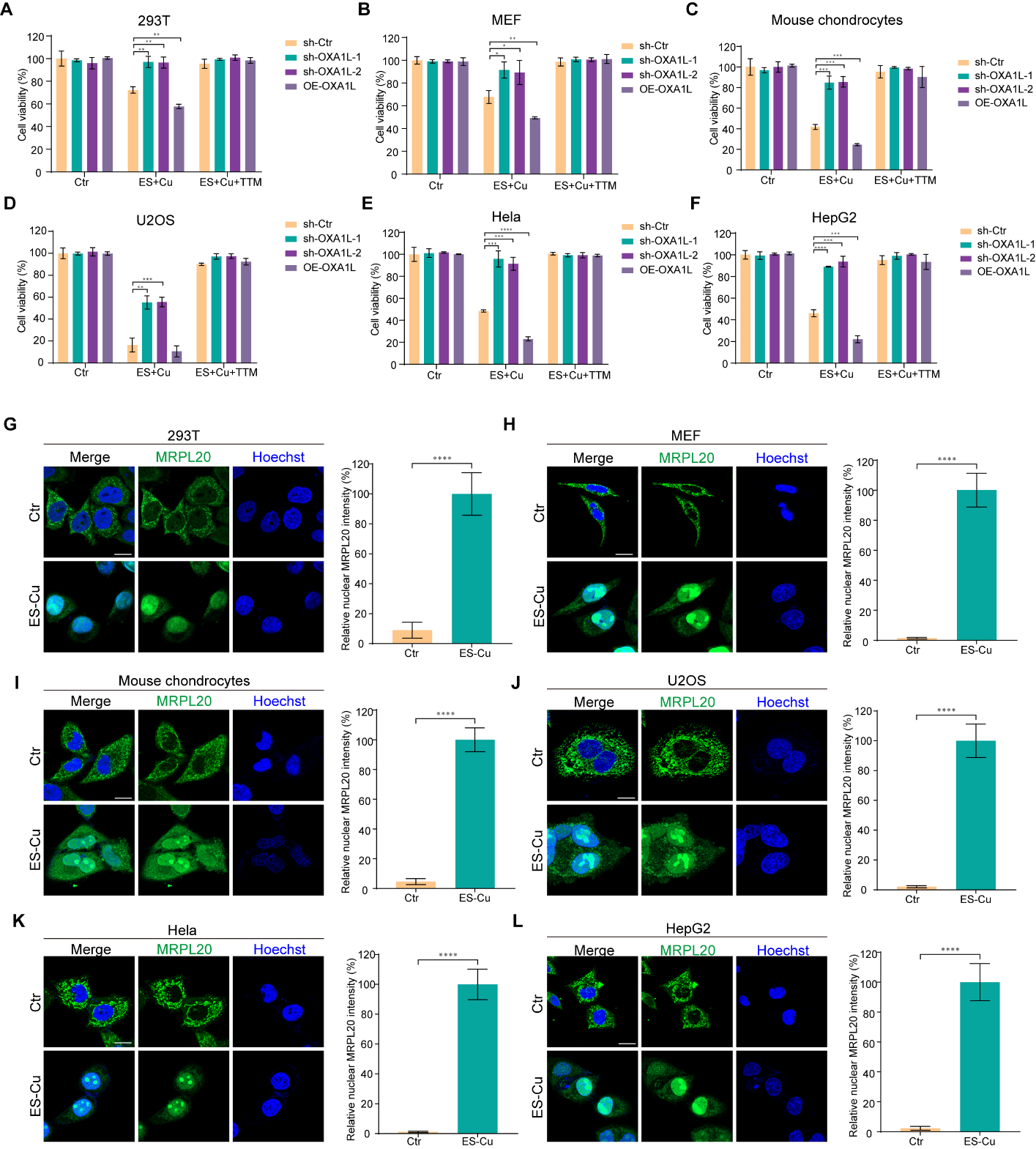

Fig. S5. OXA1L depletion protects diverse cell types from copper-induced cytotoxicity, accompanied by MRPL20 nuclear accumulation.

(A–F) CCK-8 viability measurements in 293T (A), mouse embryonic fibroblasts (MEFs; B), mouse chondrocytes (C), U2OS (D), HeLa (E) and HepG2 (F) cells expressing control shRNA, either of two OXA1L-targeting shRNAs or OXA1L. Cells received vehicle, ES–Cu (1.5 μM CuSO₄ plus 15 nM elesclomol, 10 h) or ES–Cu plus 20 μM TTM. Viability is normalized to control shRNA under vehicle conditions; mean ± SD, n = 3. (G–L) Representative images and quantification of endogenous MRPL20 (green) and Hoechst-stained nuclei (blue) in the respective cell types under control or ES–Cu conditions. Scale bars, 10 μm. Significance symbols and comparisons are shown in the panels; significance was assessed by one-way ANOVA.

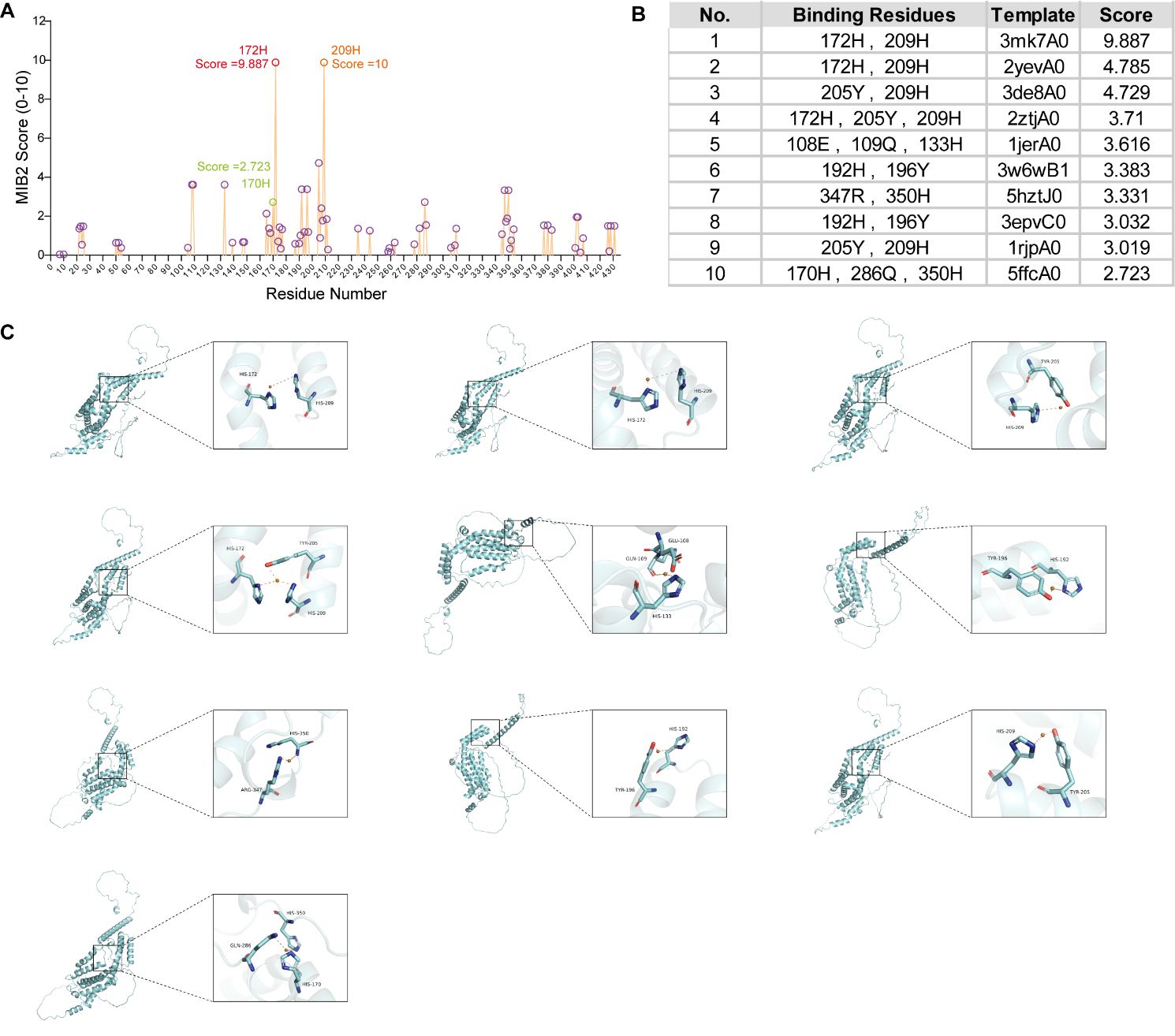

Fig. S6. MIB2 predicts candidate copper-binding sites in OXA1L.

(A) Per-residue copper-binding propensity scores for human OXA1L from MIB2; higher values indicate stronger predicted binding. (B) Ten highest-ranked metal-binding templates and the corresponding predicted OXA1L residues and scores. (C) Structural views of the ten predicted copper-binding configurations listed in (B), with enlargements of the candidate coordinating residues. These computational predictions were tested experimentally as described in the main text.

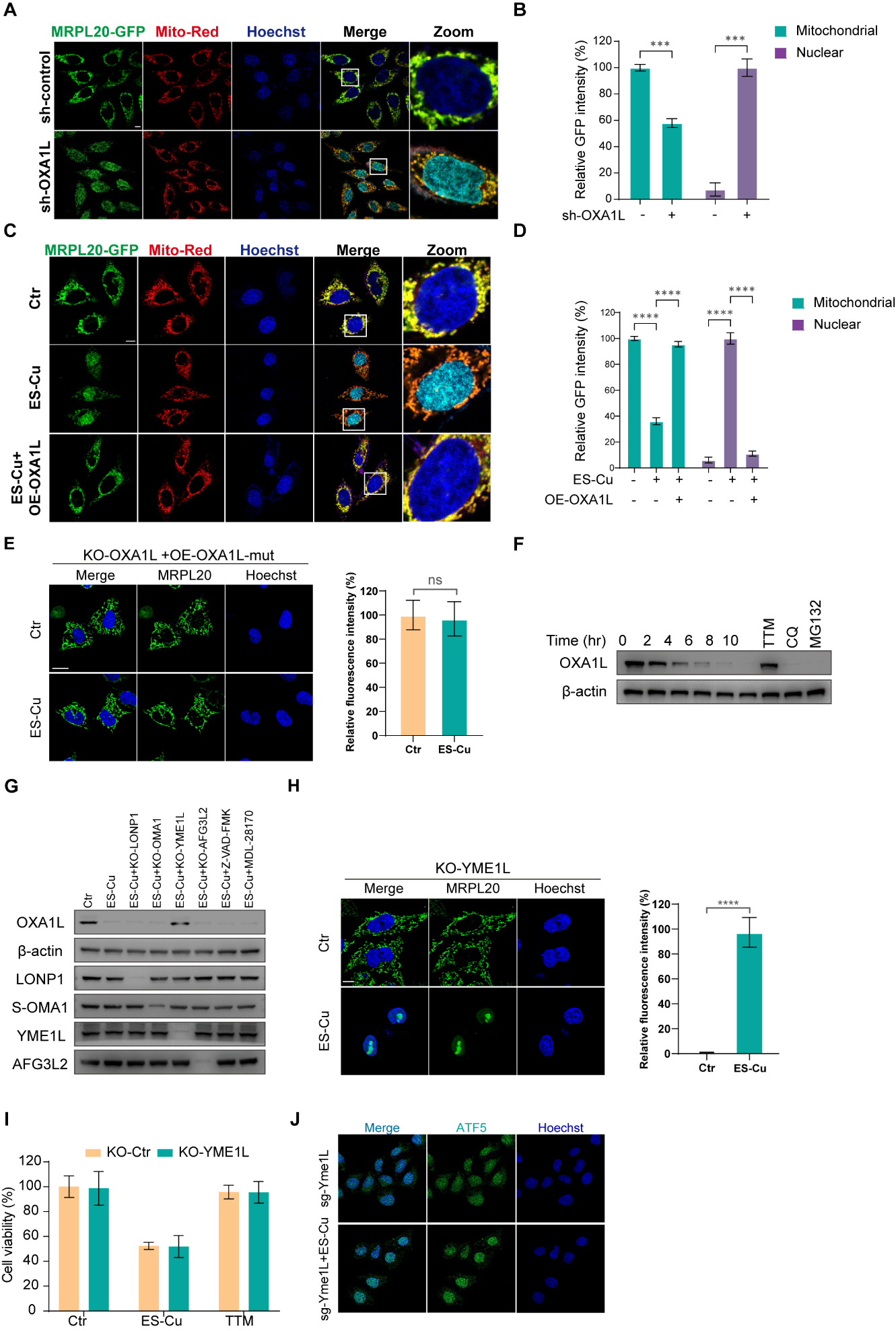

Fig. S7. OXA1L function regulates MRPL20 and ATF5 redistribution during copper stress.

(A, B) Representative MRPL20–GFP (green), Mito-Red (red) and Hoechst (blue) images in control or OXA1L-knockdown HT-1080 cells (A), with quantification of mitochondrial and nuclear GFP fluorescence (B). (C, D) Corresponding images (C) and quantification (D) from control, ES–Cu and ES–Cu plus OXA1L-overexpression conditions (1.5 μM CuSO₄ plus 15 nM elesclomol). Boxed nuclear regions are enlarged at right; scale bars, 10 μm. Quantification in (B) and (D) is mean ± SEM with n = 3 and n = 4, respectively. (E) Endogenous MRPL20 immunofluorescence (green) and nuclear fluorescence quantification in OXA1L-knockout cells expressing copper-binding-deficient OXA1L(H170A/H172A/H209A), with or without ES–Cu (10 h); Hoechst marks nuclei. (F) Time-course anti-OXA1L immunoblot following ES–Cu, with additional ES–Cu plus TTM, chloroquine (CQ) or MG132 conditions as labeled; β-actin is the loading control. (G) Immunoblots of OXA1L, LONP1, OMA1, YME1L and AFG3L2 under the indicated gene knockout conditions and ES–Cu treatment. (H) Images and quantification of endogenous MRPL20 nuclear signal in YME1L-knockout cells with or without ES–Cu (10 h). (I) Cell viability in control and YME1L-knockout cells under the indicated control, ES–Cu and TTM conditions. (J) ATF5 immunofluorescence (green) and Hoechst-stained nuclei (blue) in control and ES–Cu-treated cells carrying YME1L-targeting sgRNA. Statistical comparisons are indicated in the panels.

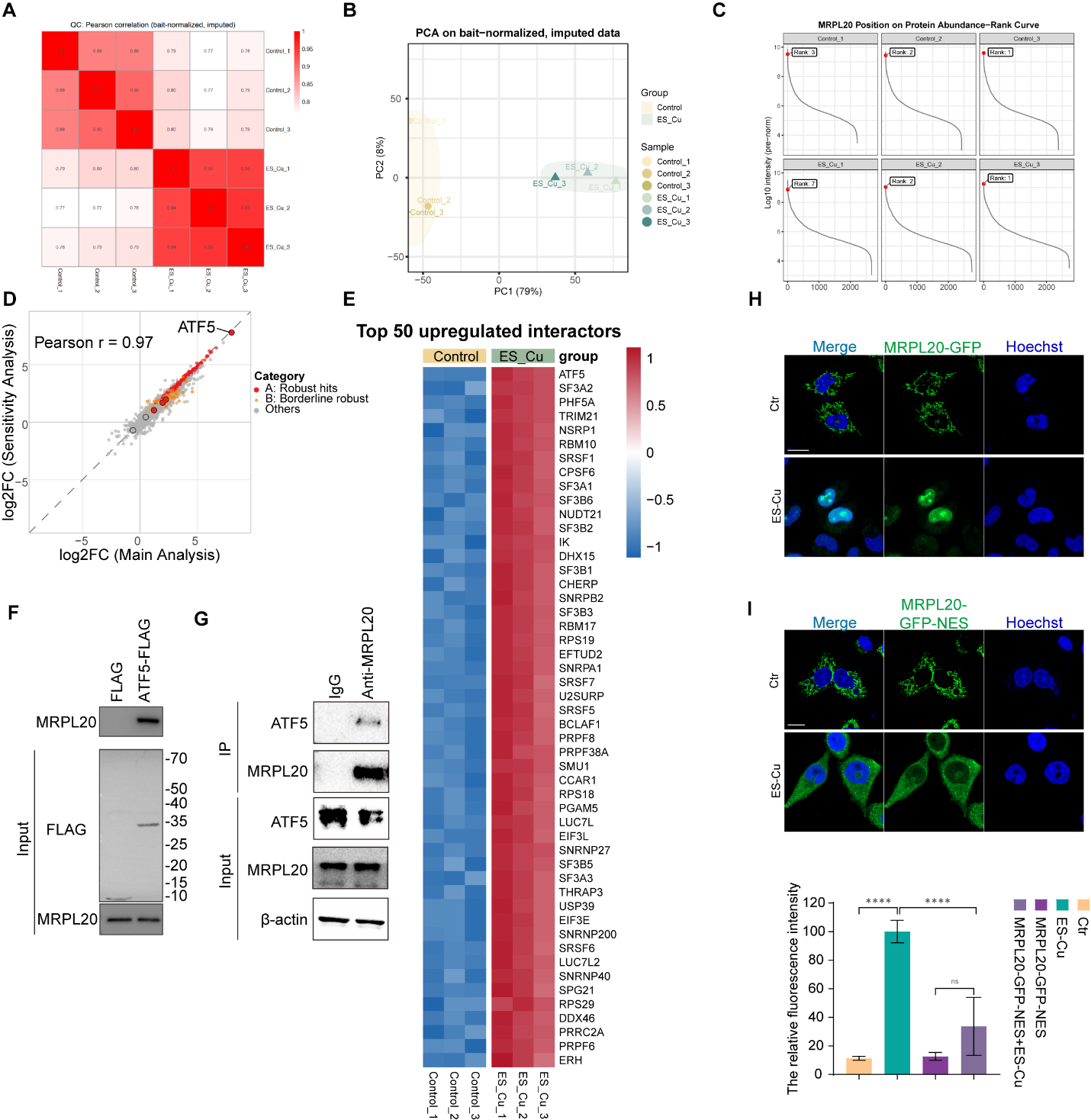

Fig. S8. Proteomic and biochemical analysis of the MRPL20–ATF5 interaction and MRPL20 nuclear targeting.

(A) Pearson correlation heatmap of the nuclear MRPL20 immunoprecipitation–mass spectrometry (IP–MS) samples (three per group). (B) Principal component analysis of bait-normalized, imputed IP–MS protein intensities in control and ES–Cu-treated groups. (C) Protein abundance–rank plots for the six IP–MS samples; red points mark bait MRPL20. (D) Comparison of interactor log₂ fold changes in the full analysis versus a sensitivity analysis excluding the ES–Cu_1 sample (Pearson r = 0.97); ATF5 is highlighted. (E) Heatmap of 50 interactions enriched after ES–Cu. (F) FLAG immunoprecipitation from cells expressing control FLAG or ATF5–FLAG followed by MRPL20 immunoblotting; inputs are shown. (G) Endogenous MRPL20 immunoprecipitation compared with an IgG control, with co-precipitated ATF5 and inputs indicated. (H) Representative images of MRPL20–GFP (green) in HT-1080 cells with or without ES–Cu (1.5 μM CuSO₄ plus 15 nM elesclomol, 10 h). (I) Representative images of MRPL20–GFP–NES (green) under the same conditions, with quantification of nuclear GFP signal for both constructs. Nuclei were counterstained with Hoechst (blue); scale bar, 10 μm. Nuclear signal is shown as mean ± SD for at least ten cells from three independent experiments; significance was assessed by one-way ANOVA.

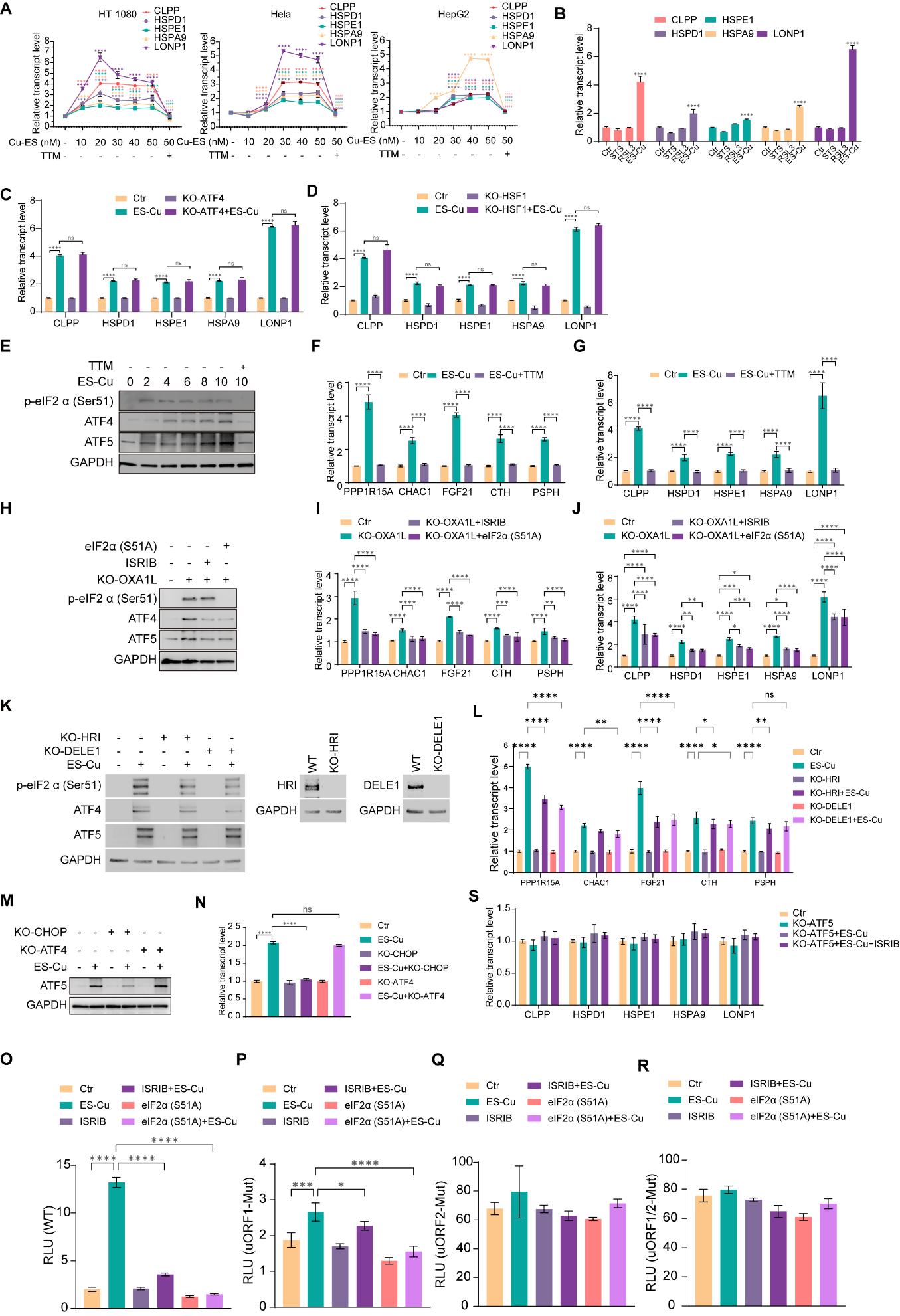

Fig. S9. Copper stress activates integrated stress signaling and ATF5-dependent UPRmt transcription.

(A) qPCR of CLPP, HSPD1, HSPE1, HSPA9 and LONP1 in HT-1080, HeLa and HepG2 cells treated with the indicated elesclomol concentrations plus CuSO₄, with a TTM rescue condition at the highest dose. (B) qPCR comparison of UPRmt gene expression after vehicle, staurosporine (STS), RSL3 or ES–Cu. (C, D) UPRmt gene expression under ES–Cu in control and ATF4-knockout (C) or HSF1-knockout (D) cells. (E) Immunoblots of phospho-eIF2α (Ser51), ATF4 and ATF5 during an ES–Cu time course, with TTM at the indicated endpoint. (F, G) qPCR of ISR transcripts PPP1R15A, CHAC1, FGF21, CTH and PSPH (F) and UPRmt transcripts (G) in control, ES–Cu and ES–Cu plus TTM conditions. (H) Immunoblots of phospho-eIF2α, ATF4 and ATF5 in control or OXA1L-knockout cells treated with ISRIB or expressing eIF2α(S51A). (I, J) ISR (I) and UPRmt (J) transcript levels for the conditions in (H). (K) Immunoblots of ISR proteins in control, HRI-knockout and DELE1-knockout cells with or without ES–Cu; HRI and DELE1 knockout validation is shown at right. (L) ISR transcript levels for the conditions in (K). (M) Immunoblot of ATF5 after CHOP or ATF4 knockout with or without ES–Cu. (N) ATF5 transcript abundance for the conditions in (M). (O–R) Reporter luminescence under control, ES–Cu, ISRIB and eIF2α(S51A) conditions for the wild-type ATF5 5′ leader (O), the uORF1 mutant (P), the uORF2 mutant (Q) and the uORF1/uORF2 double mutant (R). (S) UPRmt gene expression in ATF5-knockout cells under the indicated ES–Cu and ISRIB conditions. Error bars and statistical comparisons are shown in the panels.

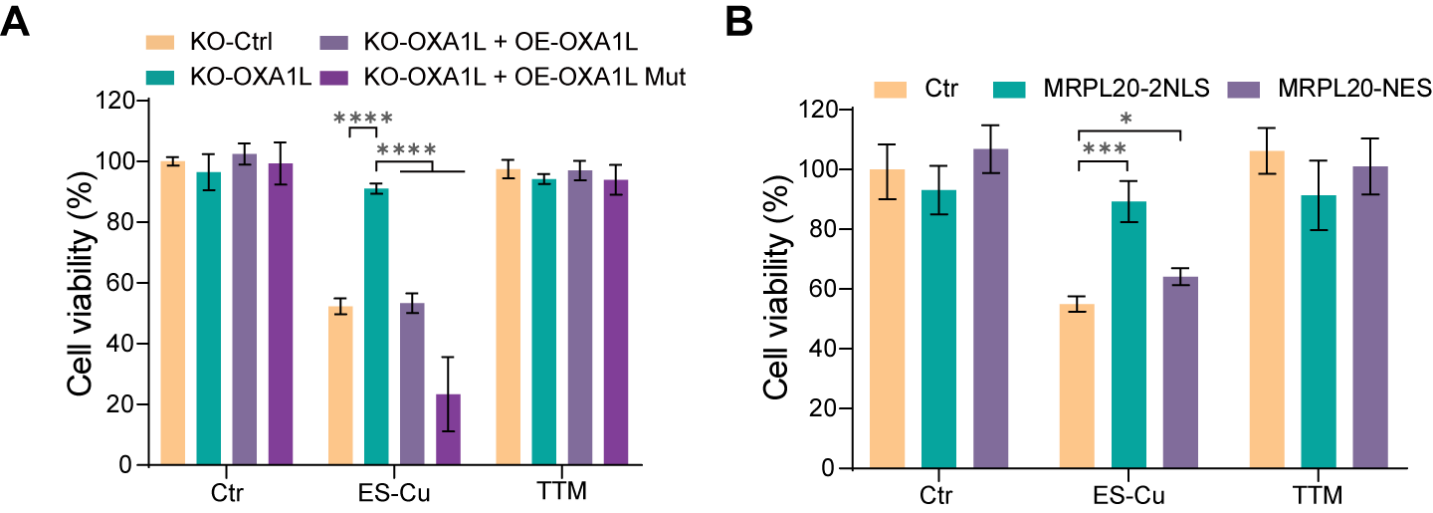

Fig. S10. OXA1L status and MRPL20 nuclear localization modulate copper-induced loss of cell viability.

(A) Viability under control, ES–Cu and TTM conditions in control cells, OXA1L-knockout cells, and OXA1L-knockout cells reconstituted with wild-type OXA1L or copper-binding-deficient OXA1L(H170A/H172A/H209A). (B) Viability under the same indicated treatments in control cells and cells expressing nuclear-targeted MRPL20–2NLS or nuclear-excluded MRPL20–NES. Group identities, error bars and statistical comparisons are indicated in the panels.

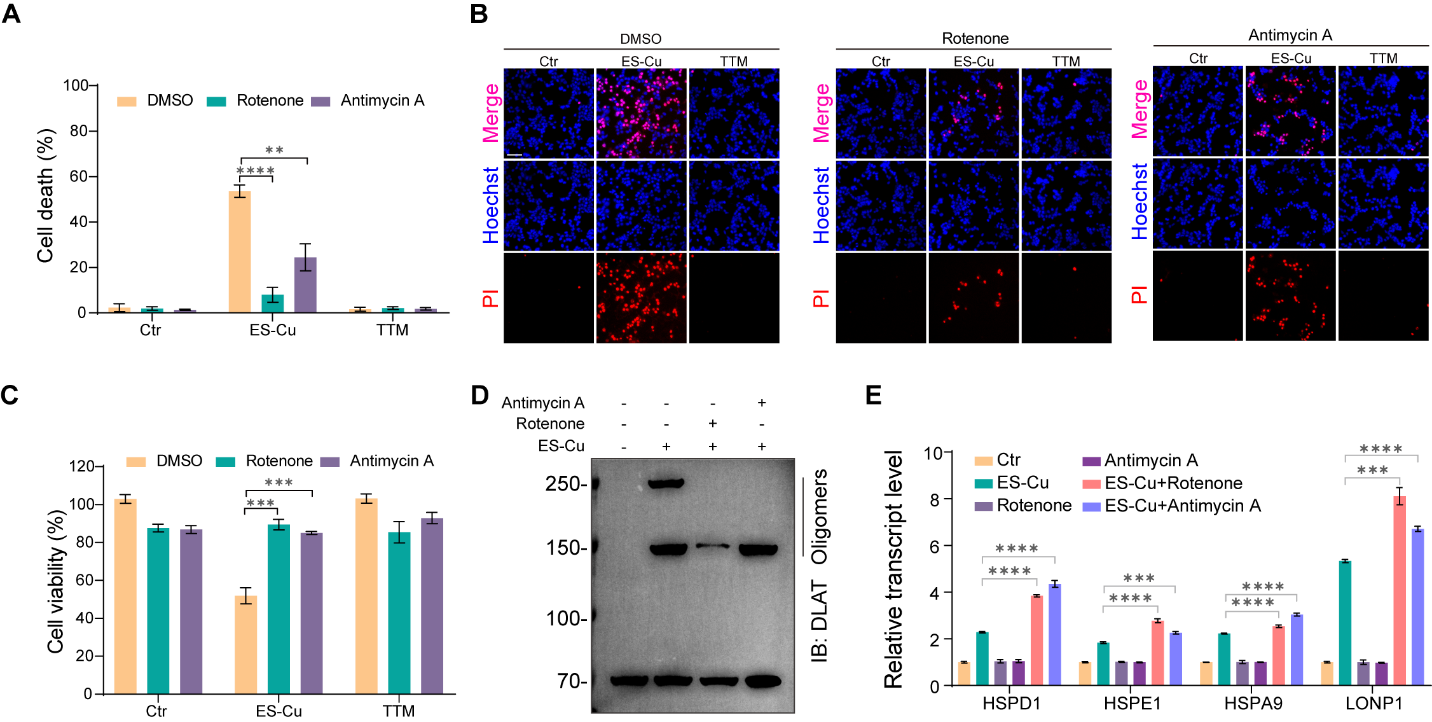

Fig. S11. Rotenone and antimycin A reduce copper-induced cytotoxicity and DLAT oligomer accumulation.

(A) PI-positive cell death under control, ES–Cu and TTM conditions with vehicle, rotenone or antimycin A. (B) Representative Hoechst (blue) and PI (red) fluorescence images of the indicated treatments. (C) CCK-8 cell viability for the treatment combinations in (A). (D) Anti-DLAT immunoblot of high-molecular-weight oligomeric species in control and ES–Cu-treated cells with or without rotenone or antimycin A. (E) qPCR of HSPD1, HSPE1, HSPA9 and LONP1 in control, ES–Cu, rotenone, antimycin A and indicated combination-treatment groups. Error bars and statistical comparisons are displayed in the panels.

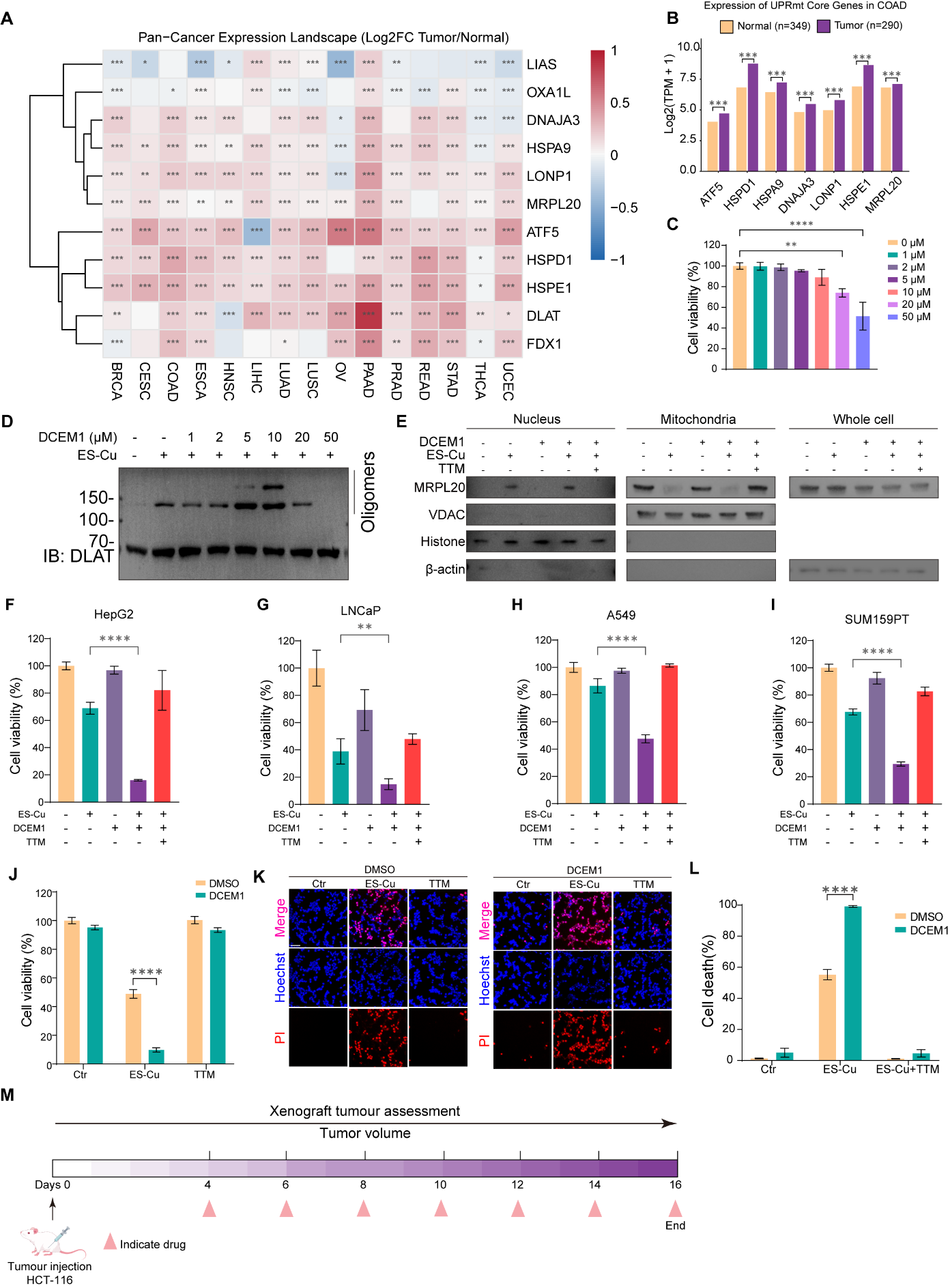

Fig. S12. Tumour UPRmt expression and copper-dependent sensitisation by DCEM1.

(A) Pan-cancer expression of UPRmt-associated and cuproptosis-related transcripts across 15 tumour types using GEPIA3 data (TCGA tumours versus TCGA and GTEx normal tissues). Color indicates log₂ fold change (tumour/normal); significance symbols are displayed in the heatmap. (B) Median expression of ATF5, HSPD1, HSPA9, DNAJA3, LONP1, HSPE1 and MRPL20 in colon adenocarcinoma versus normal tissue, shown as log₂(TPM + 1); group sizes are shown in the panel. (C) Viability of HT-1080 cells following 10 h of treatment with the indicated DCEM1 concentrations alone. (D) Anti-DLAT immunoblot of oligomeric species in HT-1080 cells treated with ES-Cu (1.5 μM CuSO₄ plus 15 nM elesclomol, 10 h) and the indicated DCEM1 concentrations. (E) MRPL20 immunoblots of nuclear and mitochondrial fractions and whole-cell lysates under the indicated ES-Cu, DCEM1 and TTM conditions. VDAC, histone H3 and β-actin serve as fractionation and loading controls. (F–I) Cell viability of HepG2 (F), LNCaP (G), A549 (H) and SUM159PT (I) cells under the indicated ES-Cu, DCEM1 and TTM combinations. (J) Viability of HCT-116 cells under control, ES-Cu or ES-Cu plus TTM conditions, with DMSO or DCEM1 as indicated. (K) Representative Hoechst (blue) and propidium iodide (PI; red) fluorescence images of HCT-116 cells under the same conditions. (L) Quantification of PI-positive HCT-116 cell death for the conditions shown in (K). The label ‘TTM’ in (J) and (K) denotes ES-Cu plus TTM. (M) Schematic of the HCT-116 xenograft experiment: tumour inoculation at day 0, administration of the indicated agents every other day from day 4 through day 16, and serial tumour volume measurements. Combination-treatment doses were ES (20 mg/kg), CuSO₄ (0.5 mg/kg), DCEM1 (20 mg/kg) and TTM (10 mg/kg).
